# Cryo-EM structure of human Nup155 reveals the biochemical basis for atrial fibrillation linked genetic mutation R391H

**DOI:** 10.1101/2021.10.05.463194

**Authors:** Sangeeta Niranjan, Jyotsana Singh, Radha Chauhan

## Abstract

Human nuclear pore complexes are composed of ∼32 distinct nucleoporins to facilitate bidirectional nucleo-cytoplasmic transport. Many of them have been associated with various human diseases such as an inherited mutation (R391H) in Nup155 is shown as the clinical cause of atrial fibrillation and sudden cardiac arrest. Due to the lack of structural knowledge and mechanistic insights, the roles of Nups in NPC assembly and relevance in human diseases are very restricted. Here, we show the cryo-EM structure of human Nup155 at 5.2-5.7. Å resolution deciphered from 3 distinct particle classes: N-terminus (19-863), C-terminus (864-1337), and longer N-terminus (19-1069). It revealed intrinsic plasticity at the middle domain of Nup155 and the role of species-specific loop regions in an atypical 7-bladed β-propeller domain to provide a distinct interface for Nup93 and Nup35. Due to the proximity of these Nups interacting sites near the Arginine-391 position, atrial fibrillation linked genetic mutation (R391H) causes dissociation from NPC in absence of N-terminal 112 residues.

**Highlights:**

- Cryo-EM structure of human Nup155 at 5.2 Å resolution
- Seven bladed β-propeller domain at N-terminus of Nup155 exhibited distinct features for interaction with Nup35 and Nup93
- The middle domain of Nup155 is highly dynamic in nature
- Structural mapping allows mechanistic interpretation of AF linked R391H mutation

## Introduction

Nuclear pore complexes (NPC) are embedded in the nuclear envelope and mediates bidirectional nucleo-cytoplasmic transport (Fahrenkrog and Aebi, 2003). Vertebrate NPC is composed of >32 different proteins called Nups to form 110MDa multiprotein complex. (Cronshaw et al., 2002). The overall architecture of NPC can be dissected into Cytoplasmic filaments, nuclear basket, and scaffold proteins such as Nup93 subcomplex that anchors central transport channel (CTC) via Nup93 (Guan et al., 1995; Hu et al., 1996; Rout et al., 2000; Zabel et al., 1996; Sachdev et al., 2012; Sonawane et al., 2020; Stuwe et al., 2015). The CTC(Nup62, Nup54, Nup58) is critical for the transport of macromolecules across the NPC (Beck and Hurt, 2017; Bui et al., 2013; Fahrenkrog and Aebi, 2003; Frey et al., 2006; Galy et al., 2003; Grandi et al., 1993, 1995; Rout et al., 2000). In spite of functional conservation across all the eukaryotes, NPCs exhibit size differences varying from 65 MDa (yeast) to 120 MDa (vertebrates).(Lin and Hoelz, 2019; Reichelt et al., 1990; Rout and Blobel, 1993).This clearly indicates significant differences in the structural organization of NPC assembly and interactome of higher organisms. (Beck, 2004; Beck et al., 2007; Bui et al., 2013; Eibauer et al., 2015; Maimon et al., 2012; Mosalaganti et al., 2018; Nudelman et al., 2018).The additional complexity of higher eukaryotic NPC is attributable to various cellular functions as mitosis, cell differentiation, chromatin organization, and gene regulation. (Belgareh et al., 2001; Breuer and Ohkura, 2015; Khan et al., 2020; Loïodice et al., 2004; Nakano et al., 2011; Strambio-De-Castillia et al., 2010).

One of the key scaffold nucleoporin Nup155 is a member of the Nup93 subcomplex (including Nup205, Nup188, and Nup35) that along with its interacting partners plays an important role in nuclear envelope assembly and cell viability. (Busayavalasa et al., 2012; Franz et al., 2005; Hawryluk-Gara et al., 2008; Vollmer et al., 2012). In vertebrates, Nup155 and Nup93 are shown to have higher abundance with 32-48 copies per NPC. (Alber et al., 2007; Cronshaw et al., 2002). In lower eukaryotes, the homologs of Nup155 (Nup157/Nup170) interacts with linker Nups; Nup93 (Nic96), Nup35 (Nup53/Nup59) (Hawryluk-Gara et al., 2005), and transmembrane Nups, Pom121 and Ndc1.(Eisenhardt et al., 2014; Hawryluk-Gara et al., 2005; Mitchell et al., 2010; Stavru et al., 2006 Onischenko et al., 2009). Due to the presence of paralogs of Nup155 (Nup157/Nup170) and Nup35 (Nup53/Nup59), the NPC composition and organization are likely to be distinct from lower to higher eukaryotes. Additionally, the low sequence conservation coverage and divergent NPC evolution (Chopra et al., 2019) further complicate extrapolating interacting regions from lower eukaryotes to higher organisms.

The structural information on Nup155 and its interacting partners is limited. The available crystal structure from lower organisms ScNup157(70-893) reveals a seven-bladed β-propeller and a downstream helical domain (Seo et al., 2013), whereas *Sc*Nup170(979-1502) shows α-solenoid domain (Whittle and Schwartz, 2009). Similarly, the crystal structure of N- and C-terminus domains from *Ct:* Nup170(74-827)•Nup53(329-361), Nup170(851-1402)•Nup145N(729-750) and Nup170(574-1402) showed conservation of N-terminal seven-bladed β-propeller domain and C-terminal solenoid domain (Lin et al., 2016). Moreover, cryo-Electron tomography-based studies on human NPC revealed ∼23 Å resolution elongated/rod shape of Nup155 that is positioned near Nup205/Nup188 (Von Appen et al., 2015). This study also revealed a membrane-binding loop (258-267) that is shown to anchor *Xenopus laevis (Xl)* Nup155 to the nuclear membrane, which seems to be conserved in yeast/*Ct*. Although these crystal structures enabled researchers to identify the basic domain organization of human Nup155. However, due to the presence of significant insertions and deletions in sequences, it is difficult to elaborate vertebrate Nup155 structure and its interaction with other members of NPC. Several multi-pronged approaches have been employed to decipher atomic resolution of the Nups and their interactions such as employing homologous thermophilic fungus structural information (Amlacher et al., 2011) coupled to crosslinking mass spectroscopy (Thierbach et al., 2013), but due to technical limitations in isolating vertebrate NPC components and their complexes, the atomic structure and interaction network of these 32 nucleoporins within NPC remains elusive. In summary, compared to the wealth of structural information available on yeast/Chaetomium Nup170/157p and its interaction with Nup53 and Nic96, our understanding of human Nup155 is poorly understood.

Several Nups are associated with various diseases such as genetic disorders, cancer, neuronal, cardiovascular, etc arising due to perturbations in NPC architecture and composition. Interestingly, Nup155 is linked with cardiac diseases such as atrial fibrillation (AF) (Leonard et al., 2020; Zhang et al., 2008) besides other reported causes of AF (Andersen et al., 2020; Parvez and Darbar, 2011a). The inherited mutation in Nup155(R391H) is shown to be the clinical cause of sudden cardiac arrest in infants. It is also shown that the R391H mutation affects nuclear envelope permeability and NPC localization (Schwartz et al., 2015; Zhang et al., 2008). However, the biochemical consequences of Nup155(R391H) mutation and how it affects NPC organization are not clear.

Here we report biochemical isolation of *Hs*Nup155 and its cryo-EM structure at 5.2 Å resolution by applying cryo-EM single particle analysis (SPA) approaches on three distinct particle classes: N-terminus (19-863), C-terminus (864-1337), and the longer Nup155 (19-1069). Our study demonstrates intrinsic plasticity in the middle domain (864-1069) of Nup155. Further, it revealed human Nup155 has vertebrate-specific insertions and deletions that provide distinct features to its β-propeller domain. We deciphered precise Nup155 interacting sites with Nup93 and Nup35 using CoRNeA (Chopra et al., 2020). Based on these interactions and structural analysis, we show that vertebrate-specific unique insertions in the 1-112 region of Nup155 are critical for Nup35 and Nup93 interaction and located in the vicinity of AF-linked genetic mutation site (R391). We also demonstrate that Nup155(R391H) mutation destabilizes its interaction with NPC due to the proximity of interacting sites for Nup93 and Nup35 present in its N-terminal 1-112 region. Altogether, our study provides an important framework for the mechanistic understanding of the role of Nup155 in NPC organization and the biochemical consequences of AF-linked R391H genetic mutation.

## Results

The multiple sequence alignment of Nup155 revealed low sequence conservation showing about 23% identity, 35% similarity with ∼82% coverage with lower eukaryotes such as *Sc* and *Ct* (Figure S1). In spite of the divergence in sequence evolution, the MSA showed overall conservation of β-propeller domain at N terminus (1-515) and α-solenoid domain at C-terminus (516-1391)(Figure S1).

### Purification of *Hs*Nup155

The details of overexpression and purification of the full-length human Nup155 are presented in the methods section. We used GFP and its nanobody as an affinity tool to purify the protein as it results in a better quality of purified protein and yield when compared with conventional affinity tags such as Ni-NTA and GST (data not shown). *Hs*Nup155 contains an amphipathic loop (258-267; known as ALPS motif) which is reported to have liposome binding activity (Von Appen et al., 2015), but the deletion of ALPS motif from *Hs*Nup155 alone didn’t solubilize the protein (data not shown) and therefore, we optimized its extraction in various detergent containing buffers (Figure 1A) and found that DDM, which is already known for its vast use in structural studies of membrane proteins (Mio and Sato, 2018; Sjöstrand et al., 2017) was able to extract Nup155 in its stable form. The final purified protein (nanobody•GFP-Nup155 complex) when analyzed by SEC-MALS and SDS-PAGE showed a homogeneous monomeric state (Figure 1B).

**Figure 1:**
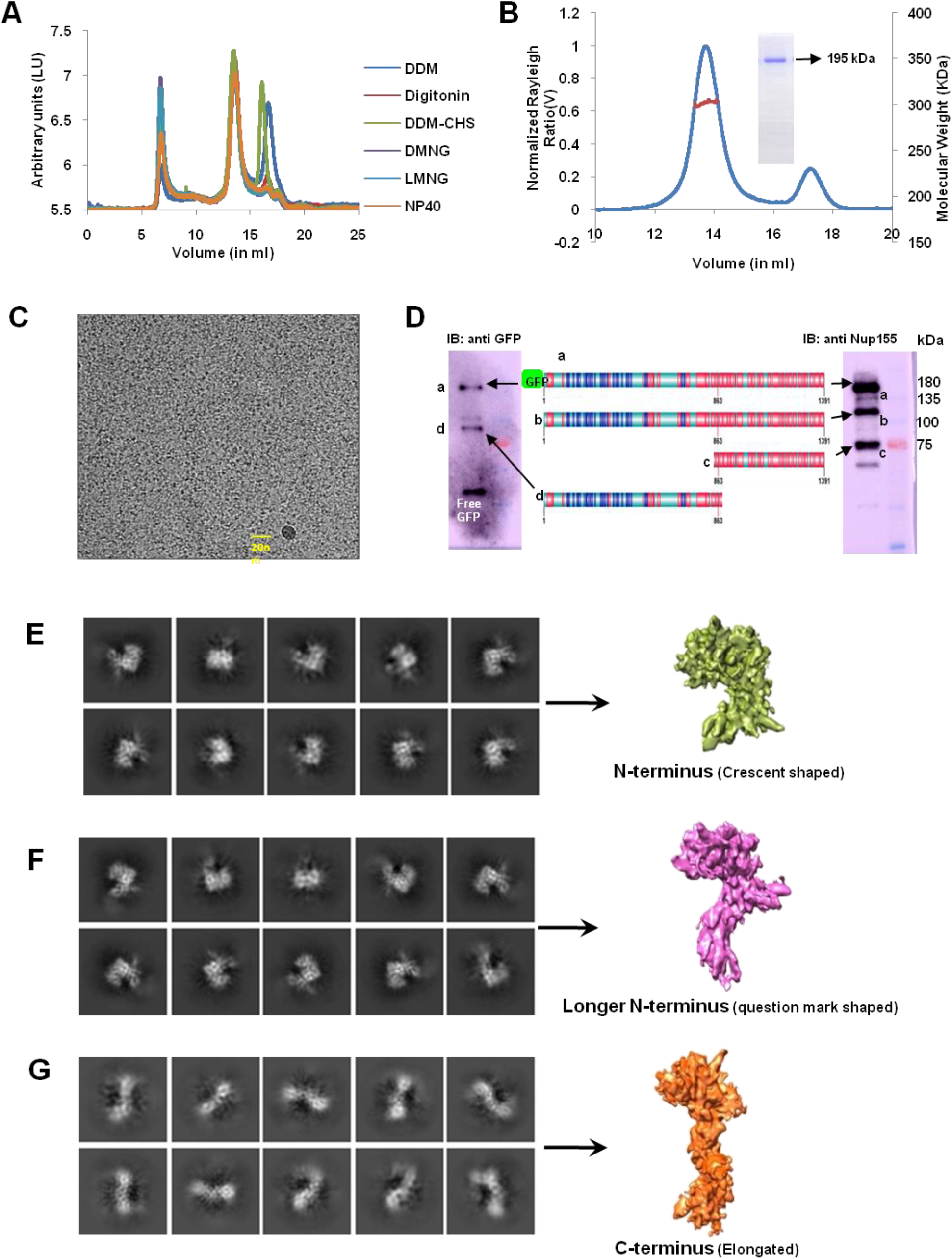
*Hs*Nup155 purification, cryo-EM sample preparation, and particle classification. (A) FSEC profile showing solubilization of GFP-fused *Hs*Nup155(1-1391) in various detergents. (B) SEC-MALS profile of purified *Hs*Nup155 and coomassie-stained gel scan of the peak fraction. (C) Representative micrograph of *Hs*Nup155. (D) Western blot of stored *Hs*Nup155 probed with anti-GFP (left) and anti-Nup155 Ab (right). Fragments indicate **a**-FL Nup155 with GFP tag(183kDa), **b**-FL Nup155 without GFP tag(155kDa), **c**-C-terminus domain (57kDa) and **d**-N-terminus domain (93kDa) with a schematic representation of *Hs*Nup155 showing secondary structures: red (α-helices), blue (β-sheets) and cyan (coiled-coil). (E-G) Representative 2D class averages for the classified particles into (E) N-terminus (green) (F) Longer N-terminus (pink), and (G) C-terminus (orange).

### Cryo-EM structure determination of *Hs*Nup155

Details of cryo grid preparation and cryo-EM data acquisition are available in Methods, Figure S2, and Table 1. We observed significant heterogeneity of purified *Hs*Nup155 protein on cryo grids (Figure 1C). About 713700 particles were auto-picked and subjected to reference-free 2D classification that revealed that the sample indeed is a mixture of at least two distinct shaped (crescent and elongated) particles. To understand the basis of this heterogeneity, we performed western blot analysis (Figure 1D) of the stored protein using anti-GFP antibody and anti-Nup155 antibody. In the case of anti-GFP immunoblot, we observed mainly 3 types of bands at different sizes: band ‘a’ (∼183 kDa) and ‘d’ (∼120 kDa) corresponding to GFP fused full-length Nup155 and N-terminal fragment, respectively. The third band corresponds to free GFP (∼27kDa). Commercially available polyclonal antiNup155 antibody is specific for its C-terminus domain (900-1050 residues), we observed 3 bands in case of antiNup155 immunoblot as well; assigned as bands ‘a’, ‘b’, and ‘c’ with the mass of ∼183,∼155 and ∼57kDa respectively). Based on their molecular size, we concluded that the band ‘a’ and ‘b’ corresponds to GFP tagged and untagged full-length Nup155 and the band ‘c’ is ∼57 kDa fragment of Nup155 corresponding to its C-terminus region. With this indication of the possible sizes from western analysis (Figure 1D) and shapes derived from homology modeling of the protein fragments (data not shown), we have concluded that the cryo-EM grid is populated with 3 major categories of particles; full-length Nup155 (with and without GFP tag), N-terminus fragment and C-terminus fragment. A rigorous 2D classification (Figure S2) of all particles resulted into 3 categories of 2D classes (Fig 1E-G) referring to N-terminus (crescent-shaped: category a; Figure 1E), longer N-terminus/FL (question-mark shaped: category b; Figure 1F), and C-terminus (elongated particles: category c; Figure 1G). Henceforth, each category was independently subjected to 2D and 3D classification and was processed further for de novo structure determination (Figure S2 and Methods).

**Table 1.** Cryo-EM Data Collection, Refinements and Validation Statistics

| Data Collection and Processing |  |  |  |
| --- | --- | --- | --- |
| Voltage (kV) | 300 |  |  |
| Electron exposure<br>(e-/Å<sup>2</sup>) | 57.6 |  |  |
| Defocus range (µm) | -1.0 to -2.5 |  |  |
| Pixel Size (Å) | 0.823 |  |  |
| Symmetry imposed | no |  |  |
| Initial Particle images (No.) | 713700 |  |  |
|  | HsNup155<br>N-ter | HsNup155<br>Longer N-ter | HsNup155<br>C-ter |
| Final Particle Images | 88600 | 85000 | 78600 |
| Map resolution (Å) (FSC<br>threshold) | 5.2 | 5.7 | 5.3 |
| Refinement |  |  |  |
| Initial Model usedfor<br>homology modelling (PDB<br>ID) | 5HAX | 5HAX and 5HB1 | 5HB1 and 3I5P |
| Map Sharpening<br>B factor (Å<sup>2</sup>) | -460 | -475 | -535 |
| Model Composition |  |  |  |
| Protein Residues | (19-863) | (19-1069) | (864-1337) |
| Ligands | 0 | 0 | 0 |
| Root-Mean-Square-Deviations |  |  |  |
| Bond lengths (Å) | 0.008 | 0.007 | 0.007 |
| Bond angles (°) | 1.244 | 1.365 | 1.249 |
| Validation |  |  |  |
| MolProbity score | 2.62 | 2.85 | 2.63 |
| Clashscore | 51.96 | 85.97 | 45.07 |
| Poor rotamers(%) | 1.52 | 0.7 | 0.7 |
| Ramachandran Plot |  |  |  |
| Favored (%) | 95.75 | 93.12 | 91.53 |
| Allowed (%) | 3.81 | 6.27 | 7.20 |
| Disallowed (%) | 0.44 | 0.61 | 1.27 |

The final EM maps when subjected to the local resolution analysis showed a uniform range of resolution spread in N-terminus (Figure S3A), C-terminus (Figure S3D), and longer N-terminus (Figure S3G). We obtained a final GSFSC resolution of 5.2 Å, 5.3Å, and 5.7Å for N-terminus (Figure S3B), C-terminus (Figure S3E), and longer N-terminus (Figure S3H) respectively as well as good coverage for orientations (Figure S3C, S3F, and S3I respectively). Overall, the electron density maps were true to their GSFSC resolution and showed continuous well-ordered density for N-terminus (Figure 2), C-terminus (Figure 3), and longer N-terminus (Figure 4). We noted well-resolved densities for helices (Figure S4A, Figure 2B, Figure 3B) and β-propeller blades (Figure S4B) as expected for ∼5-5.5Å resolution (Casañal et al., 2019). The presence of good density regions for few helices such as C704-Q726 of N-terminus (Figure S4A) and S1237-Y1255 of C-terminus (Figure 3C) as well as loop region (Figure 3D) showed confidence in sequence assignment for model building, for which, initial homology models were edited based on the respective EM density maps. The final refined model showed overall good geometry (Table 1) and we could build 3D models from 19-863 in N-terminus (Figure 2A, 2B); 864-1337 for C-terminus (Figure 3A, 3B), and 19-1069 for longer N-terminus structures (Figure 4B, 4C and Table 1). Notably, the electron density maps of the N-terminus (Figure S4C) and the C-terminus region (Figure 3E) clearly revealed that the Nup155 fragmentation occurred at (863-864) residue that is in agreement with our western blot analysis (Figure 1D).

**Figure 2:**
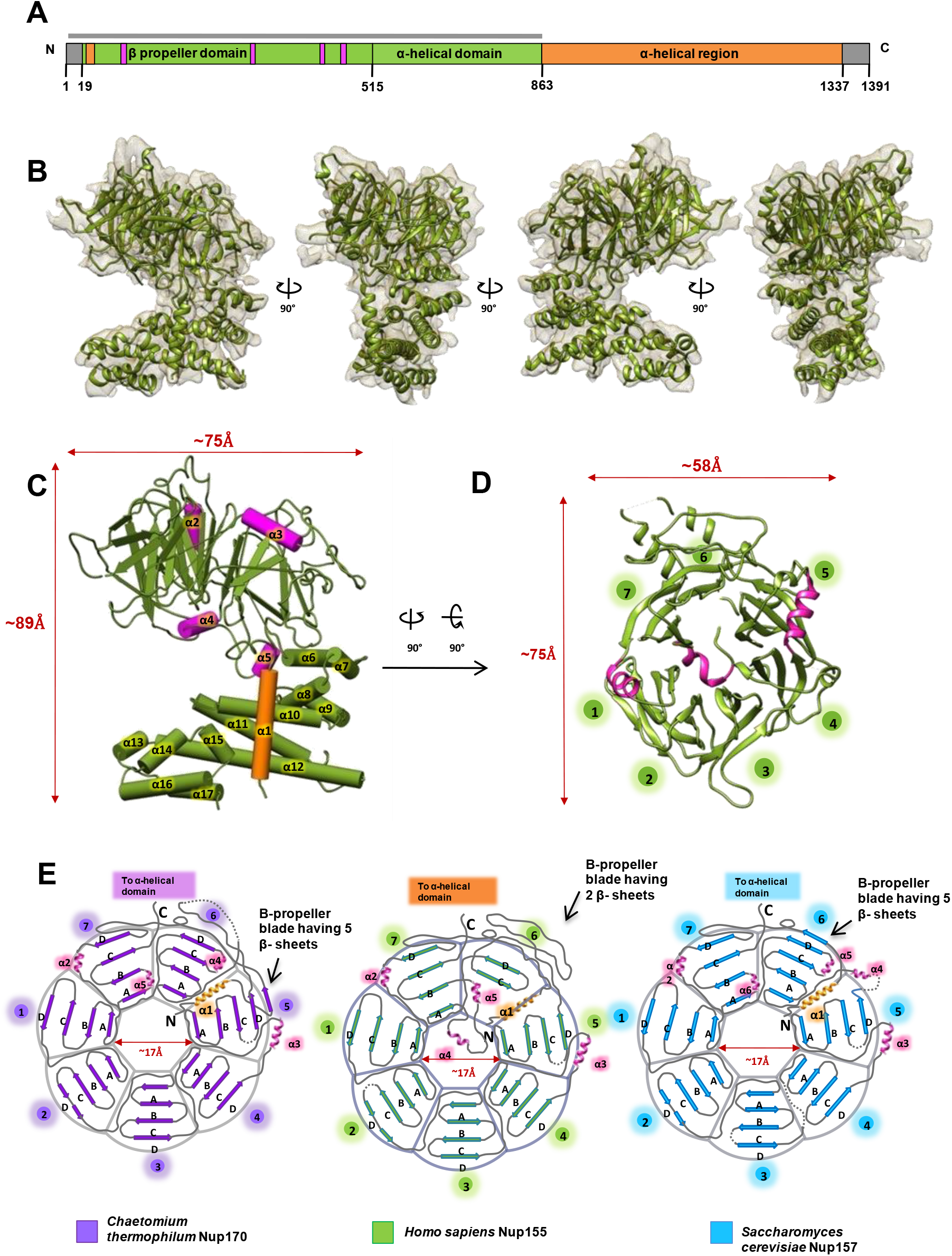
3D Structural details of *Hs*Nup155 (19-863) N-terminus domain. (A) Schematic representation of domain architecture of *Hs*Nup155. The bar above the domain architecture corresponds to the solved Cryo-EM structure of the N-terminus region (19-863). (B) Ribbon representation of N-terminus structure fitted into Cryo-electron density map (5.2Å) shown at 90° rotation angle. (C) Schematic representation of N-terminus domain starting with the first helix (orange), β-propeller domain (green) with interrupted helices α2 to α5(pink) and α-helices from α5 to α17 (green). (D) The ribbon representation of β-barrel structure showing the arrangement of β-sheets forming 7-bladed β-barrel and location of its α-helical elements, color scheme same as in ‘C’. (E) Schematic representation of β-barrel domains of *Ct, Hs*, and *Sc* and location of its individual β-sheets and its differential composition of the β-sheets to form the 7-bladed arrangement of the barrel and intermittent α-helical elements. Dotted gray lines-disordered regions.

**Figure 3:**
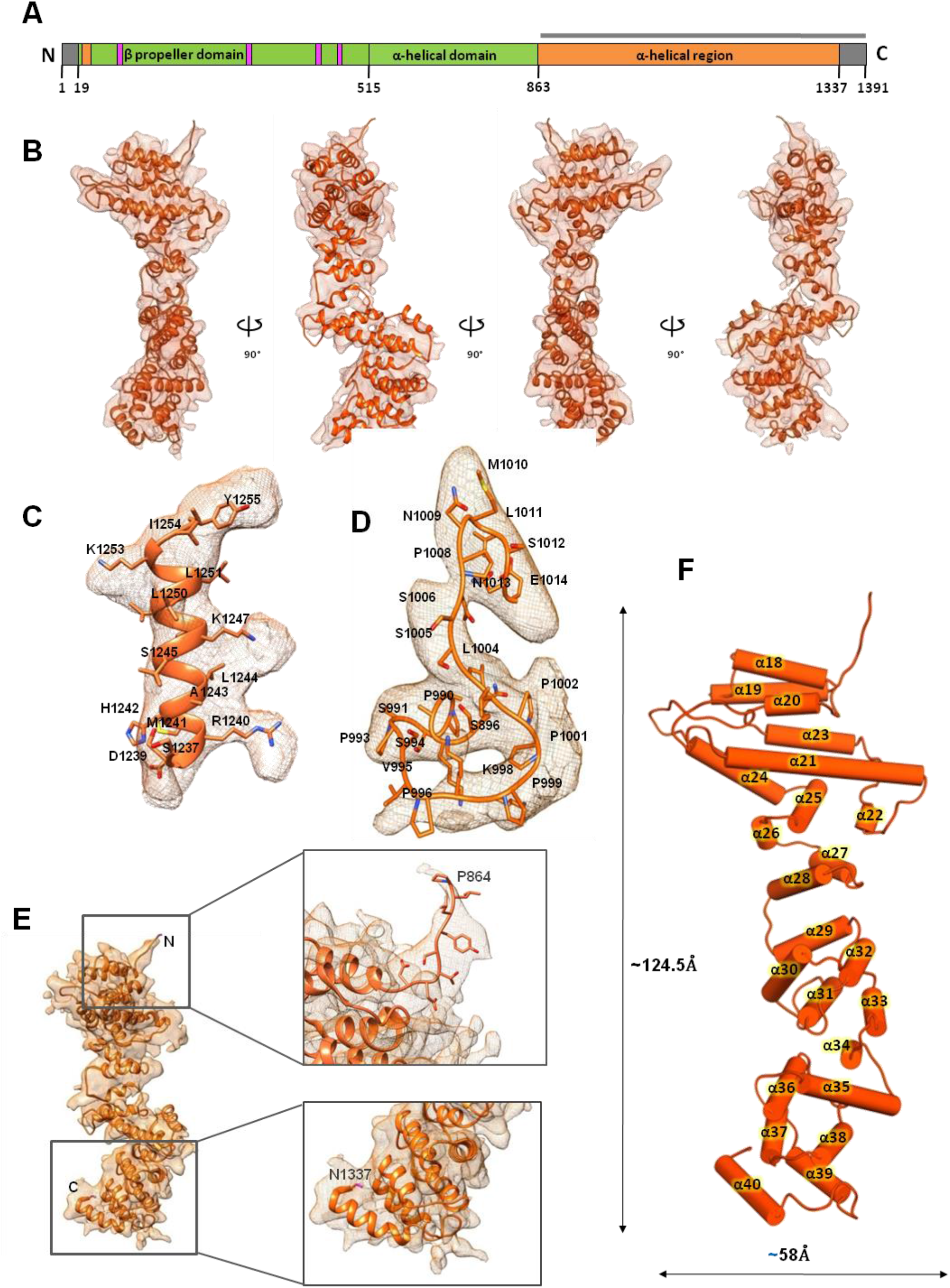
3D Structural details of *Hs*Nup155 (864-1337) C-terminus domain. (A) Schematic representation of domain architecture of *Hs*Nup155. The bar above the domain architecture corresponds to the Cryo-EM structure of the C-terminus region(864-1337). (B) Ribbon representation of C-terminus structure fitted into Cryo-electron density map (5.3Å) shown at 90° rotation angles. (C and D) CryoEM density maps and fits for helix α36 and interhelix loop between α23 and α24 (985-1014) respectively (E). The density coverage of the C-terminus domain showing terminal residues: 864 and 1337. (F) Cartoon representation of the arrangement of helices in the C-terminal domain numbered from α18 to α40 along with dimensions.

**Figure 4:**
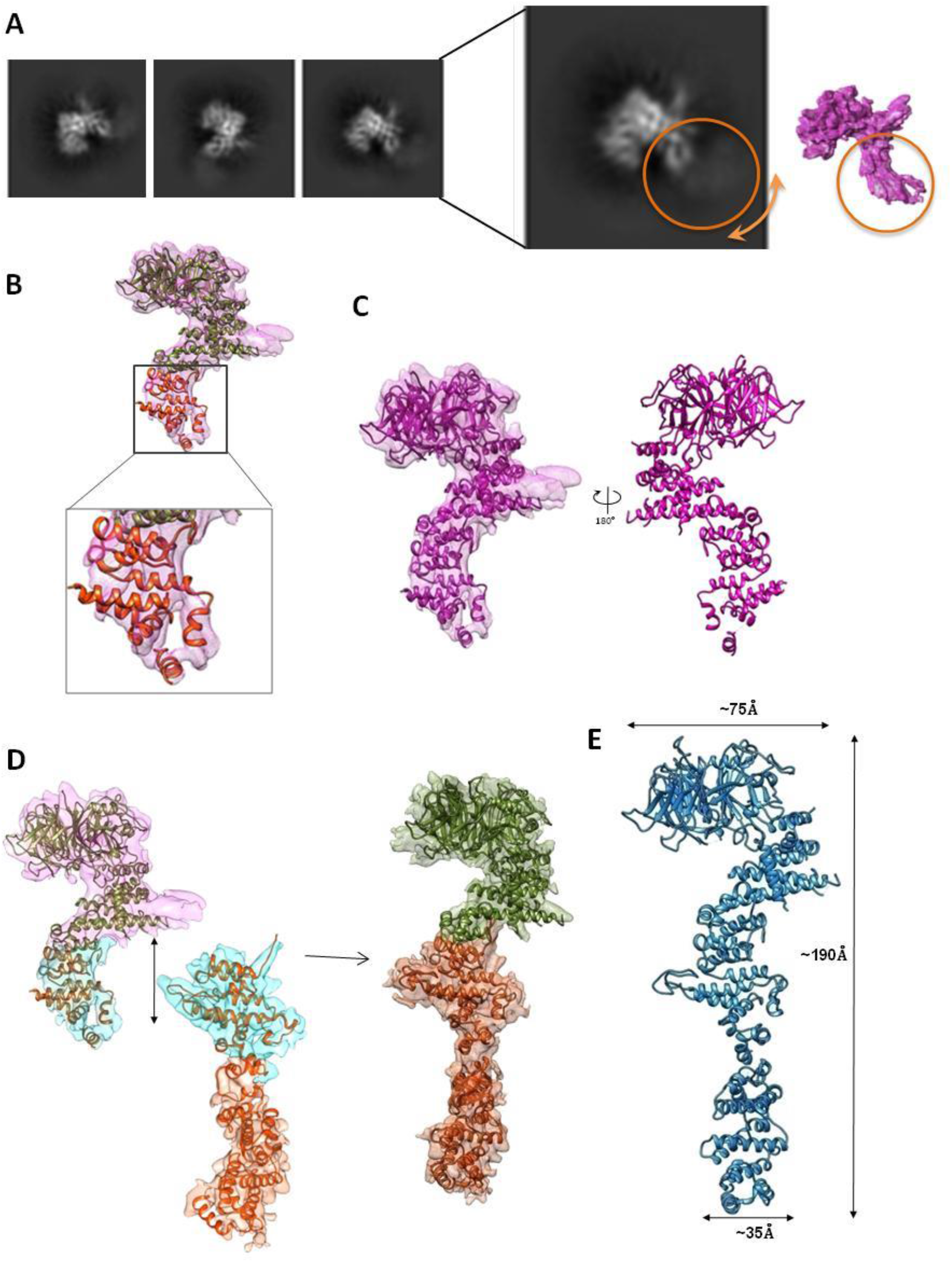
Structural analysis of Longer N-terminus density map. (A) Representative 2D classes from Category B particles (longer N-terminus; question-mark shaped) and its corresponding 3D map (pink). The ‘fuzzy tail’ characteristic highlighted using orange circle (B) Structure of *Hs*Nup155 (19-1069 residues; pink) is elucidated by placing *Hs*Nup155 N-terminus structure (19-863; green) and further building downstream (864-1069) at C-terminal region based on density map. (C) Ribbon representation of the structure of Longer N-terminus (19-1069; pink) fitted into corresponding EM density map (left) and its 90-degree view (right) (D) The overlapping region (cyan) in longer N-terminus and C-terminus structure are superposed to create a full-length model of *Hs*Nup155. (E) Cartoon representation of the full-length model of Nup155 (19-1337) and its dimensions.

### The cryo-EM structure of *Hs*Nup155 (19-863) region and comparison with lower eukaryotes

Nup155(19-863) is composed of β-propeller domain followed by the α-helical domain (Figure 2B) which folds to form a compact crescent-shaped structure with overall dimensions of ∼89Å × 75Å × 58Å (Figures 2C and 2D) and the final model is refined at GSFSC_0.143_; resolution 5.2Å. The N-terminal domain starts with the first α-helix (Fig 2C; orange-colored α1), which is located perpendicular to the plane of the β-propeller domain and continues to form a solenoid structure. This β-propeller domain is composed of radially arranged 7 blades of β-strands (Figure 2D). These blades are formed by anti-parallel connected strands starting from innermost strands arising outwards and forming individual blades arranged around a circular funnel shaped-pore (Figure 2D). The outermost strand of the 7^th^ blade (7D) is the first β-sheet from the N-terminus and it interacts with the corresponding inner strand (7C) to complete the four-stranded 7^th^ blade. The α-helices extend from the end of the 7th blade of the β-propeller domain and continue to form the alpha-solenoid domain in a zigzag trend (Fig 2C and 2D). A few small α-helical insertions (α2, α3, and α4) in between β-sheets help in forming the interface between β-propeller blades and α-helical domain via α5. α2 and α3 are positioned in 7D1A and 4D5A interblade loops of β-propeller respectively, whereas α4 and α5 both are positioned in the same interstrand loop at 7AB (Figure 2D and 2E). The positioning of α1, α4, and α5 provides the interface between β-propeller and α-helical domains (Fig 2C). 7C is the last strand of the β-propeller domain which exits to begin the formation of α-helical domain with α6 and α7 helices situated anti-parallel to each other. These α-helices continue to grow into an elongated α-solenoid domain containing α6 to α17 helices. We did not observe any electron density for few disordered regions; initial 1-18 aa, 185-192aa in 2CD loop and 377-381aa in 5CD loop of the β-propeller domain, 530-531aa, and 579-658aa which includes the longest insertion region (585-646) of 61 residues found uniquely in vertebrates and located between α9–α10 helices, 680-701aa located between α10–α11 helices, 734-765aa located between α11–α12 helices (Fig S1).

The homologous structures of the Nup155 N-terminus domain are available from *Sc* (PDB: 4MHC and 3I5P) (Seo et al., 2013; Whittle and Schwartz, 2009) and *Ct* (PDB: 5HAX and 5HB1) (Lin et al., 2016). For structural comparison, Nup155 and its homologous structures are superpositioned (Cα RMSD of 4.453 and 6.358 with PDB: 5HAX and 4MHC respectively), (Figure S5A). We noticed that the canonical seven-bladed β-propeller domain is highly conserved among lower eukaryotes (Lin et al., 2016; Seo et al., 2013), but there are substantial differences in their sequence and structural composition of vertebrate Nup155 (Figure S1, Figure S5A). The 6^th^ blade in *Hs*Nup155 is atypically composed of only 2 β-strands (named as blades 6C and 6D) while *Ct*Nup170 has canonical 4 β-strands (named as blades 6A to 6D) in its 6^th^ blade (Figure 2D and 2E). The corresponding region in *Ct*Nup170, which forms β-sheets 6A and 6B (in continuation of blade 6C), is a deletion in the sequence of *Hs*Nup155 and thereby, we found no density for the missing 6A and 6B blades (Figure S4F). *Sc*Nup157 and *Ct*Nup170 have 5 β-strands in their 6^th^ and 5^th^ blades respectively (Figure 2E). The α2 and α3 helices are also conserved across these species while α4 helix is unique to *Hs*Nup155 positioned with a conserved α5 helix (α6 in *Sc*), across these species (Figure 2E). The α4 helix is an insertion unique to *Hs*Nup155 and is located at the interface of the β-propeller and α-solenoid domain. Therefore, when superpositioned, the corresponding region is found to be missing in *Ct*Nup170 (Figure S4D). The position of the α4 helix is also found to create some unique interactions with interblade loops of the β-propeller domain. Residue I473 forms direct interaction with residues D222 and N286 (Figure S4E). Such evolutionarily distinct changes suggest the emerging differences in canonically conserved β-barrels, which are primarily evolving due to various insertions and deletions in sequences. Altogether, these unique features of the human Nup155 7-bladed β-propeller clearly show the inhered flexibility and versatility to preserve the basic fold yet provide uniqueness. Such observation for WD40 fold is although previously reported (Afanasieva et al., 2019), yet was an interesting observation for NPC evolution indicating that diversity in sequence and structures extend to diversity in acquiring complexity in functions.

There are two evolutionary conserved motifs present in Nup155 (Von Appen et al., 2015; Lin et al., 2016) and we observed ordered density for both motifs (Figure S5B). The first is the WF motif, composed of evolutionarily conserved (Fig S1) and solvent-exposed tryptophan and phenylalanine residues (246-247) in the 3CD loop (Fig S5B). Previously WF motif in *Ct*Nup170 is also predicted to reinforce membrane binding (Lin et al., 2016). The second motif which resides in the 3D4A loop of both the homologous protein *Hs*Nup155 and *Ct*Nup170 (Figure S1) is predicted to form an amphipathic loop characterized by the absence of charged residues; a unique feature of the ALPS motif. This loop contains a universally conserved proline residue (Figure S1; Figure S5B) on the polar face of the helix (Lin et al., 2016) along with serine and leucine residues. This loop was deleted in the case of the *Ct*Nup170 crystal structure. However, we observed a very clear electron density for the ALPS motif in the *Hs*Nup155 structure (Fig S5B and S5C).

### The Cryo-EM structure of the *Hs*Nup155 (864-1337) region

We observed well-ordered density (GSFSC_0.143_ resolution 5.3Å) for the 864-1337 region in Nup155 C-terminus EM structure (Figure 3A and 3B). However, we were not able to capture density for the last 54 residues (1338-1391), probably due to its flexible nature (Figure 5E), therefore, this region was omitted in the final model building and refinement. We observed mostly well-resolved helices and loops in the density map (Figure 3C and 3D). The final model showed that the Nup155 (864-1337) is composed of only α-helices (α18 to α40) arranged in a zigzag fashion with interhelical loops forming an elongated α-solenoid region with overall dimensions ∼125Å × 58Å × 52Å (Fig 3F). The *Hs*Nup155 (864-1337) structure also revealed the clear electron density for longer loops such as S896-E1014 (Figure 3D), which were not resolved/reported in homologous structures from *Ct*Nup170 and *Sc*Nup170p (Lin et al., 2016; Whittle and Schwartz, 2009).

**Figure 5:**
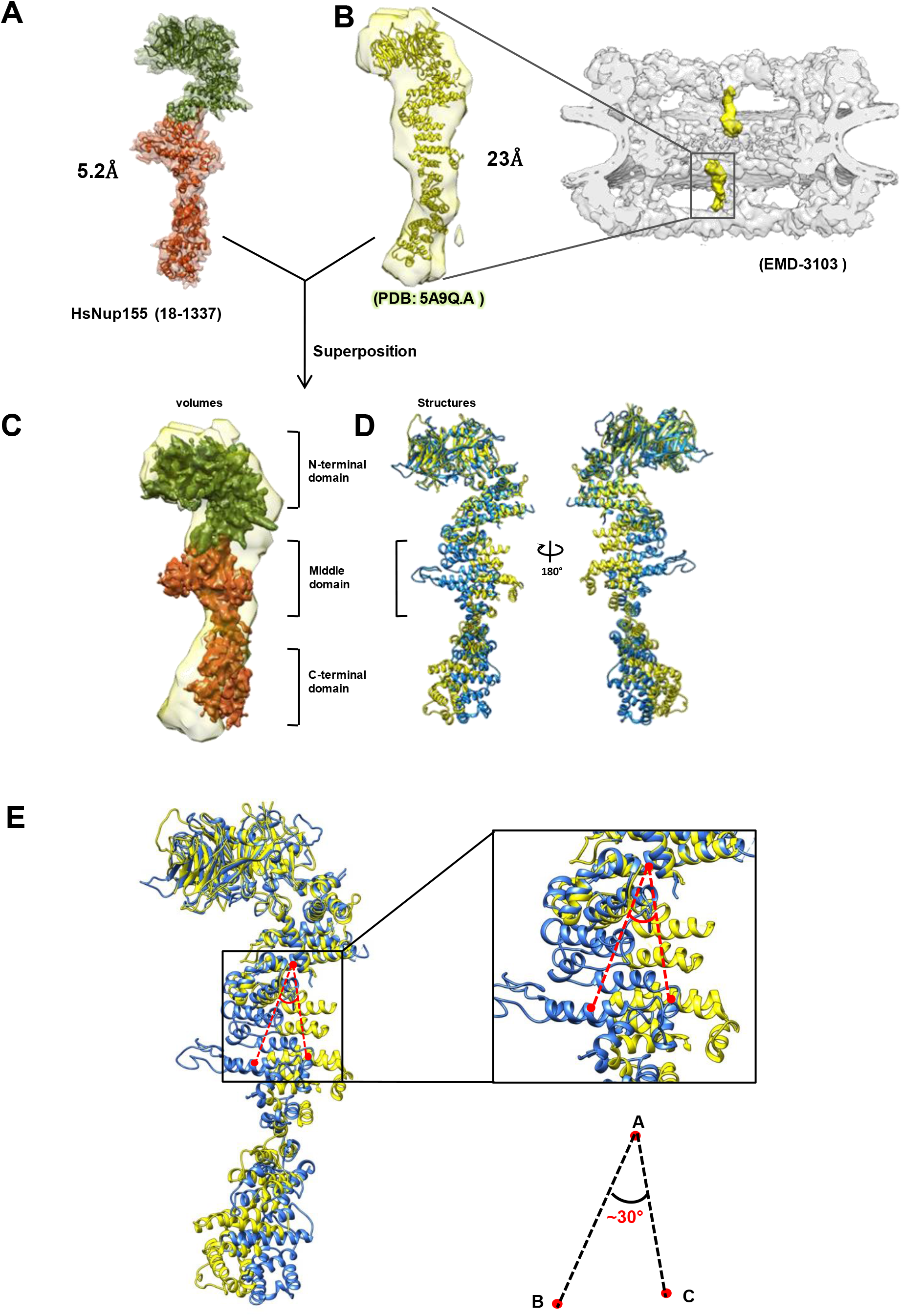
Comparison of *Hs*Nup155 (18-1337) structure with tomography density. (A) Cartoon representation of *Hs*Nup155 Cryo-EM structures fitted into CryoEM densities of Nup155 domains (N-terminus: green and C-terminus: orange) connected to form full-length Nup155 model. (B) 3D model of *Hs*Nup155 from PDB 5A9Q.A (yellow) was fitted in the respective density of Nup155 (yellow) in human cryo-ET density (EMD-3103). (C) Superposition of Cryo-EM densities of *Hs*Nup155 N-terminus (green) and C-terminus (orange) onto tomography density map (yellow) indicating three distinct regions: N-terminus, middle and C-terminus. (D) Superposition of full-length structure of *Hs*Nup155 (light blue) with PDB:5A9Q.A. (yellow) (E) Measurement of angle for movement/displacement in the α-helical region of *Hs*Nup155 in comparison with PDB:5A9Q.A. Points A, B and C represent residues 791 (superposed in both structures), 974 (Nup155), and 974(5A9Q.A) respectively.

Upon superposition (Figure S5D) with homologous crystal structures (PDB: 5HB1 from *Ct* and PDB: 3I5P from *Sc;* Cα RMSD of 4.739 and 6.846, respectively), we observed that although the overall helical arrangement of the *Hs*Nup155 (864-1337) remain similar, there is a significant shift: particularly 864-1105 region, which does not align well (Figure S5D). Overall *Hs*Nup155 fragment formed an elongated structure and showed significant differences from homologous structures as observed in *Hs*Nup155 N-terminus (19-863) structure.

### The Cryo-EM model of *Hs*Nup155 (19-1337) based on longer N-terminus structure

We have obtained a question-mark-shaped longer N-terminus Nup155 (19-1069) density map (Figure S2) and the 2D classes obtained clearly show the presence of crescent-shape at one end indicating the presence of β-propeller domain that is continuous to fuzzy-cloud characteristics at another end suggesting the possible flexibility in this region (Figure 4A). Upon 3D reconstruction, we obtained a well resolved EM density map (Figure S3G, Figure 4B) refined to GSFSC_0.143_ resolution of 5.7Å (Figure S3H) with decent particles orientation coverage (Figure S3I) resulting in reasonably resolved helices (Figure S4G). Due to its exhibited flexible nature at its C-terminus (Figure 4A), we could model only the 18-1069 region of the Nup155 with good geometry and refinement parameters (Table 1; Figure 4C). We noted that the β-propeller domain followed by the solenoid domain was almost identical to the Nup155 (19-864) structure (Cα RMSD of 0.273).

Nup155 (19-1069) structure showed that helix α17 is extended further to form compact domain 864-1069. This region is overlapping with the structure of *Hs*Nup155 (864-1337) and aligns very well. We, therefore, have utilized this longer N-terminus structure to build the full-length model of *Hs*Nup155_19-1337_ by overlapping the 864-1069 domain from the longer N-terminus with Nup155 (864-1337) structure (Figure 4D). This composite structure that covers from 19-1337 region revealed a longitudinal dimension of ∼190Å (Figure 4E) indicating a highly elongated shape for *Hs*Nup155.

The tomography-based map of human NPC at a resolution of 23Å is available (Kosinski et al., 2016). To validate our composite (19-1337) structure, we performed both density fit and superposition with the reported model of Nup155 (PDB: 5A9Q.A; (Von Appen et al., 2015)). We observed that our density map (Figure 5A) is fitting well in 5A9Q: A (Figure 5B and 5C). When we superposed our cryo-EM model of *Hs*Nup155 (19-1337) with PDB: 5A9Q.A chain homology model revealed Cα RMSD of 9.6A with a major shift in Nup155 middle (864-1105) and C-terminal end (1110-1337); Figure 5D). We further calculated the possible angle of movement in 864-1069 α-helical domain using point ‘A’ marked on overlapping 791^st^ residue while point ‘B’ and ‘C’ are non-overlapping 974^th^ residues on both the models and observed a possible ∼30° angle of movement in this middle domain of the α-helical region (Figure 5E) thus clearly demonstrating the dynamic nature or movement of Nup155 (864-1069) middle region perhaps due to its intrinsic plasticity. Such movement was evident in our 2D classes (Figure 4A), due to which we could not obtain a well-resolved density for our full-length Nup155 particles. It is noteworthy that *Hs*Nup155 C-terminus (864-1391) region is when separated; it facilitates the stability and homogeneity of the protein. Moreover, such dynamic nature of the middle domain of Nup155 is also evident in previously reported homologous crystal structures of *Ct*Nup170 (PDB:5HB0 and 5HB1) (Lin et al., 2016) indicating the evolutionary significance of such conformational changes.

### Protein-protein interaction network of Nup155 partners

Nup155 is known to co-purify with Nup93 and Nup35 (Amlacher et al., 2011; Eisenhardt et al., 2014; Hawryluk-Gara et al., 2005, 2008; Lin et al., 2016). We investigated the Nup155 interaction network using *insilico* CoRNeA method to predict interacting interfaces from primary sequences (Chopra et al., 2020). When we analyzed the top 10% of interacting pairs for Nup35•Nup155 (Figure S6A), Nup93•Nup155 (Figure S6B) and Nup93•Nup35 (Figure S6C), we observed that the highest convolution scores correspond to the N terminal (34-49) and the RRM domain (170-250) of *Hs*Nup35 to interact with the N-terminal (85-89) and C-terminal region (1087-1089) of *Hs*Nup155 respectively. (Figure 6A and S6A). The C-terminal region of *Hs*Nup35 is also predicted to interact with *Hs*Nup155, however with lower scores (Figure S6A). We further compared these predicted interfaces with lower eukaryotes known interacting regions (Lin et al., 2016) using MSA (Figure S7A), which revealed some notable differences in the interfaces such as region corresponding to *Ct*Nup53(329-388) that binds to Nup170 (Amlacher et al., 2011), is missing in vertebrates (Figure S7A). Moreover, our predicted interface region corresponding to *Xl*Nup53(162-320) that binds to Nup155 (Eisenhardt et al., 2014) is highly conserved among vertebrates. Overall, we concluded that *Hs*Nup35 is likely to pose a distinct interface for Nup155 as compared to the Nup53•Nup170 interaction interface.

**Figure 6:**
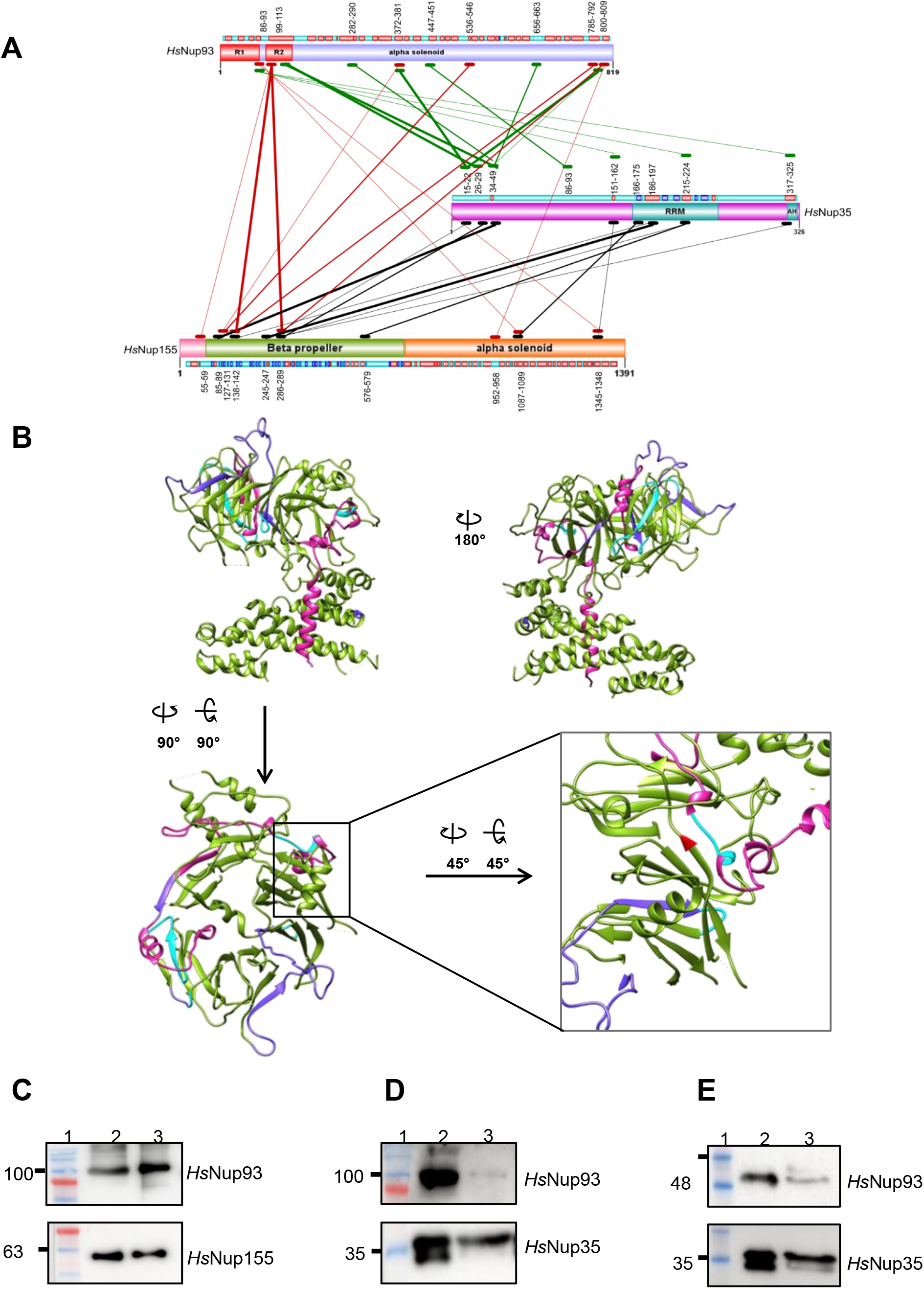
Structural mapping of CoRNeA predicted interactions of Nup155 with Nup93 and Nup35. (A) Schematic representation of top 10% the interface regions among *Hs*Nup35, *Hs*Nup93 and *Hs*Nup155. 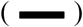; highest-scoring, 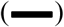; intermediate scoring and 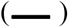; low scoring residue pairs. 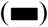; range of amino acid residues forming an interacting interface. Black, Red and green lines show predicted interacting interface between *Hs*Nup35-*Hs*Nup155, *Hs*Nup93-*Hs*Nup155, *Hs*Nup35-*Hs*Nup93 respectively. (B) Nup155 N-terminus structure (olive green) is highlighted for (19-112) residues in pink, Cornea predicted Nup93 and Nup35 interacting regions highlighted in cyan and purple, respectively. Box shows zoomed region with R391 (Red) and interacting interface with insertion 52-59, predicted to be interacting with Nup93 also (cyan). (C, D, and E) GST-pulldown assay performed with GST fused *Hs*Nup93. Pulled fractions containing His-tagged *Hs*Nup155(1-515) and *Hs*Nup35(1-305) analyzed by Western blot using Abs, Lane: 1- Marker, 2-Input, 3-Eluant.

In the case of *Hs*Nup93•*Hs*Nup35 interacting pair, the top 10% predicted interface revealed the region of Nup35 at the N-terminus (38-42) to interact with the Nup93 N-terminus (99-113) along with various middle and C-terminal regions (371-381, 796-819 and 656-663). These predictions when compared with known interaction from previous studies (Lin et al., 2016) showed that *Ct*Nup53 N-terminus (30-70) that binds to Nic96 (Amlacher et al., 2011) is poorly conserved (Figure S7B). It is also shown that *Xl*Nup35 N-terminal(84-167) is crucial for Nup93 binding (Eisenhardt et al., 2014; Hawryluk-Gara et al., 2008). Our CoRNeA analysis too predicted 86-93 regions of *Hs*Nup35 to interact with *Hs*Nup93. Similarly, in the case of *Hs*Nup93•*Hs*Nup155 interacting pair, the top 10% predicted interface showed that *Hs*Nup93(96-107) along with the C-terminal region (536-809) forms an interface with various regions of the *Hs*Nup155 N-terminus (286-289, 138-142, 127-131) located mostly in the β-propeller domain (Figure 6B). Mapping of both Nup35 and Nup93 interacting interface on Nup155 revealed interface clustering on the β-propeller domain (Figure 6B), which is in agreement with a previously reported study in *X. laevis* where it was shown that Nup93(608-820) region stabilizes the interaction of Nup35 and Nup155 (Sachdev et al., 2012). Similarly, a study in *Drosophila* also highlights the interaction of Nup155 NTD/β-propeller domain (1-539) with Nup93 (Busayavalasa et al., 2012).

In summary, we observed that Nup93, Nup35, and Nup155 have interacting sites for each other (Figure 6A) and *Hs*Nup35 harbors overlapping interfaces with both Nup155 and Nup93. It is also noteworthy that the Nup93 N-terminus coiled-coil region is known to interact with the CTC (Sonawane et al., 2020), Nup205/Nup188 (Chopra et al., 2020), and also important for interaction with Nup35 and Nup155. Such interaction is not shown previously including lower eukaryotic species. Therefore, to validate CoRNeA predictions, we co-purified His6-tagged *Hs*Nup155 (1-515) and His6-tagged *Hs*Nup35 (1-305) along with GST-tagged Nup93 (96-819) region (Figure 6C and 6D) and (1-150) region (Figure 6E) using affinity chromatography using GST tag (See methods). Western analysis of these purified complexes clearly revealed that both variants of *Hs*Nup93 (96-819) and (1-150) are capable to copurify with *Hs*Nup35 (1-305) (Figure 6D and 6E) showing predicted (96-107) region of *Hs*Nup93 is indeed a true interactor for Nup35.

### Structural basis of the AF linked genetic mutation R391H

The cryo-EM structure of *Hs*Nup155 (19-864) revealed that conserved AF-linked genetic mutation R391H (Figure 7A), is present on the 5^th^ blade of the β-propeller domain (Figure 7B). The R391 is positioned at the end of the β-sheet (384-391 on blade 5D) and is likely to not cause any structural change when mutated to Histidine. However, we noticed that there are two vertebrate-specific unique insertions in the sequences, which form loops (52-59 and 69-74) in the *Hs*Nup155 (19-864) structure. We further identified that loop 52-59 forms a direct interaction with R391 residue (Fig 7B). To find out the structural difference with lower eukaryotes, we have superposed the structures of *Ct*Nup170 (PDB: 5HAX) and *Hs*Nup155. In the case of *Ct*Nup170 the 5^th^ blade is composed of 5 β-sheets (Figure 6B). However, in the case of HsNup155, the insertion of 52-59 loop resulted in the loss of the 5th β-sheet secondary structure. We also observed that both insertions are evolutionarily conserved among vertebrates (Figure S1).

**Figure 7:**
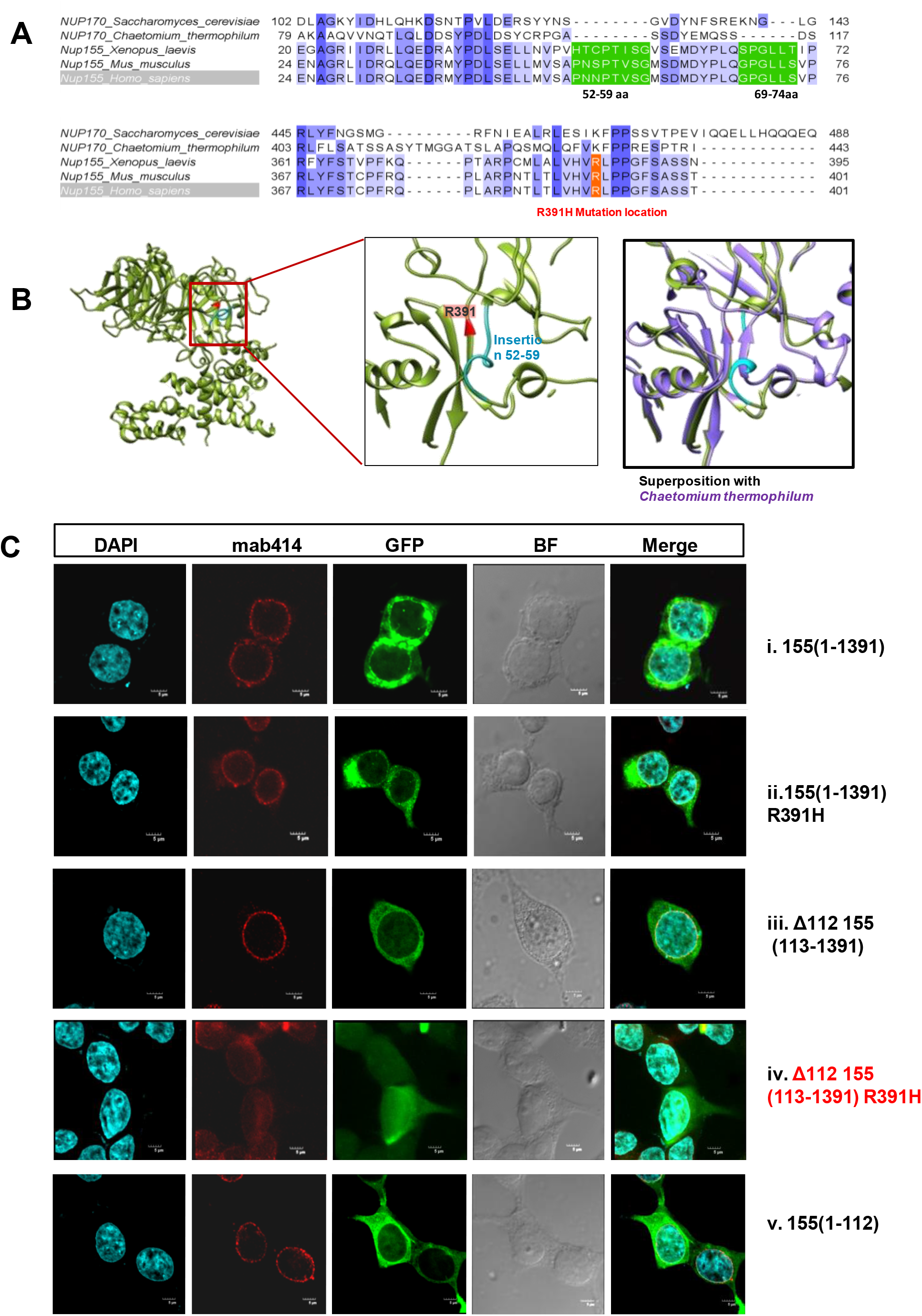
*Hs*Nup155(R391H) mutational analysis. (A) MSA of *Hs*Nup155 showing unique loops (52-59 and 69-74) conserved in vertebrates. The R391 residue is highlighted in red color. (B) Cartoon representation of *Hs*Nup155(19-863) structure showing the positioning of R391H (red) and 52-59 loop (cyan) in the vicinity. The zoomed image shows the superposition of the *Hs*Nup155 (19-863) structure (green) onto the corresponding *Ct* structure (purple) revealing the loss of the β-sheet in *Hs*Nup155(19-863) region due to 52-59 loop. (C) GFP immunofluorescence analysis of variants of *Hs*Nup155. DAPI and mab414 is used to stain nuclei (cyan) and NPC (red), respectively, GFP fused over-expressed Nup155 variants (green) are: i *Hs*Nup155 (1-1391), ii: *Hs*Nup155(1-1391)R391H and iii: *Hs*Nup155(113-1391) localizing onto NPC. iv: Nup155 (113-1391)R391H mutant showing mislocalization and v: Nup155 (1-112) alone is localizing onto NPC. The scale bar represents 5µm.

Altogether, based on our structural analysis of R391H mutation (Figure 7B) as well as interaction analysis (Figure 6A) we hypothesized the following: The N-terminus of Nup155(1-112) is critical for its interaction with its partners and the presence of two insertion loops in the proximity of R391 residue (52-59 and 69-74) would alter interaction network of R391H mutation. To validate, we performed a GFP immunofluorescence-based localization assay of Nup155 and its various variants as shown in Figures 7C and S6E. We have observed that the full-length protein, Nup155 (1-1391) is capable of localizing to the NPC, creating a nice GFP rim, while Nup155 (1-1391) (R391H) mutant showed partial loss of localization (Figure 7C). Similarly, the Nup155 (1-750) is capable of localizing to the NPC on the nuclear rim, while its mutant Nup155 (1-750) (R391H) showed partial loss of localization (Figure S6E). The truncation of the 1-112 region in both Nup155 (1-1391) and Nup155 (1-750) showed minimal loss of localization (Figure 7C and S6E). Interestingly, combining the mutation R391H with deletion of 1-112 residues, in Nup155 (1-1391) showed the localization on the nuclear envelope is drastically affected including the change in morphology of the nuclear envelope (Figure 7C). In Nup155 (1-750) also, the combination of the mutation R391H with deletion of 1-112 residues affected the localization as well as nuclear pore permeability (Figure S6E). However, Nup155 (1-112) construct exhibited its proper localization on the NPC (Figure 7C) in spite of its very small size indicating its importance for protein-protein interaction, which is in agreement with our CoRNeA-based interaction studies. Based on this, we concluded that 1-112 region of *Hs*Nup155 plays an important role in interaction with other Nups at NPC and drives the localization of Nup155 on the nuclear envelope and due to its presence in proximity of R391 site, would adversely affect the interaction network in *Hs*Nup155 (R391H) mutant.

## Discussion

Cryo-electron tomography is extensively applied to understand the overall NPC architecture in both lower and higher eukaryotes (Von Appen et al., 2015; Lin et al., 2016; Mosalaganti et al., 2018). However, the resolution is still restricted to >20 Å. The overall architecture derived from these studies along with our comparative analysis (Chopra et al., 2019) clearly underlines the species-specific Nups or their subcomplexes arrangement. In the case of human NPC, the challenges in biochemical isolation of Nups or their subcomplexes have restricted our understanding of how each Nup plays its role in the NPC assembly and transport functions. Moreover, in spite of its direct relevance in many human diseases, the underlying molecular mechanisms of these Nups remain elusive. In our present study, we show biochemical isolation of *Hs*Nup155 for structural analysis. By using the SPA based cryo-EM method, its structure is determined to ∼5-5.5Å resolution that enabled us to understand its overall structure, plasticity and biochemical consequences of AF linked R391H mutation.

The purification of *Hs*Nup155 using GFP as an affinity tag guided us to segregate the N-terminal and C-terminal particles from the heterogeneous sample. Moreover, the GFP fusion tag was beneficial for screening various detergents using FSEC and showed DDM to be suitable for extraction and cryo-EM analysis of Nup155. Similar strategies can now be applied to other larger human Nups to isolate them for structural studies and thus can lead to atomic resolution structures of human NPC or its sub-complexes.

Cryo-EM SPA approach is routinely used to obtain atomic-resolution structures of proteins and their assemblies and relies on averaging of randomly oriented protein particles. In the case of Nup155, due to partial cleavage of the protein at 863-864 residues (Figure 1D, 3E, and S4C), created three different kinds of particles having different shapes and dimensions. This condition ultimately helped us in capturing the small (57 kDa) C-terminal α-solenoid domain (864-1337) separately. It is noteworthy that in spite of the smaller size of this fragment, the structure to 5.3Å resolution could be determined in C1 symmetry indicating the stability of the C-terminus fragment when cleaved. In spite of having >50% particles of full-length protein on the grid, we were unable to capture and reach 2D classes because of the highly elongated shape (190Å×75Å×35Å) of the protein-coupled with the dynamic behavior of the middle α-solenoid domain. Moreover, current tools for particle picking are optimal for globular proteins and new strategies are required for highly elongated particles. Overall, our study demonstrates a strategy to solve the SPA-based cryo-EM structures of dynamic nucleoporins and their challenges, which will subsequently help in reaching a step closer to understanding the structure and dynamics of the NPC.

*Hs*Nup155 has an atypical β-propeller domain which is significantly distinct from its homologs in lower eukaryotes (Figure 2E). The β-propeller domain is one of the most conserved protein domains found abundantly from prokaryotes to eukaryotes (Kopec and Lupas, 2013) and is defined by the symmetric arrangement of 4-12 propeller blades (each blade formed by 4 anti-parallel β sheets) in a toroidal fashion. Among them 7-bladed β-propeller domain is suggested to be the most favored geometric arrangement (Murzin, 1992) and also associated with a broader variety of functions (Chen et al., 2011). However, in the β-propeller domain of Nup155, it is evident that vertebrates have numerous insertions and deletions leading to (Figure S1) significant deviations (such as loss of 2 β-sheets in 6^th^ blade) from the classical 7-bladed β-propeller domain (Figure S4F). Altogether, we demonstrate that the human Nup155 β-propeller domain has acquired uniqueness throughout evolution, perhaps to cater the needs of functional complexity of higher eukaryotes.

Many nuclear envelope genes including NPC components have been shown as candidates that impact normal cardiac functions (Anter et al., 2009; Han et al., 2019; Marzlin, 2020; Nanni et al., 2016; Nofrini et al., 2016; Parvez and Darbar, 2011b; Tarazón et al., 2012; Zhang et al., 2008). Particularly, more than 160 genes have been associated with Atrial Fibrillation (AF) which is multifactorial cardiovascular dysfunction leading to cardiac failure (Andersen et al., 2020). A recent study revealed that the N-terminal β-propeller domain is the hotspot, where most of the clinically linked allelic variants of NUP155 (24 variants) are clustered and present around R391H and L503F mutations (Leonard et al., 2020; Zhang et al., 2016, 2008). However, it was not clear how this β-propeller domain is affected by these variants. Our study based on Nup155(19-863) structure, interaction, and immunofluorescence analysis revealed the presence of unique insertions (52-59 and 69-74) in close proximity of β-sheet containing R391 (Figure 7B), that directly interacts with R391th residue and the initial 100 residues of Nup155 are significant to interact with Nup35 and Nup93. We conclude that the Nup155 allelic variant hotspot is also a hub for other Nups interactions and thus would destabilize their interactions. Such insights would be immensely beneficial in postulating a testable hypothesis to dissect the role of Nup155 in cardiac development or disorders.

It is previously reported that the composition and stoichiometry of the NPC vary with tissues (Ori et al., 2013). However, it is not clear how compositional NPC variations are acquired by NPC in different tissues. Interestingly, there are three isoforms reported for *Hs*Nup155 in which isoform2 (1-59 residues missing) and isoform3 (812-868 missing) are splice variants of isoform1 (1-1391). Nup155 isoform2 lacks the initial residues from 1-59, thereby missing the unique insertion loop for interaction with R391. Since the effect of R391H mutation is linked to cardiac cells, we propose that Nup155 isoform2 might be more prevalent than isoform1. Moreover, isoform3 is lacking 812-868 residues as compared to isoform1 and thereby missing 3α-helices (α15-α17), which will probably result in the reduced longitudinal dimension of Nup155. We propose that these different isoforms might play significant roles in cell/tissue-based specificity and development.

Taken together, our study for the first time demonstrates human Nup155 structure and its comparison across species revealed significant differences particularly in interacting interfaces with Nup35 and Nup93. The approach to use single-particle analysis using Cryo-EM is promising for the study of dynamic nucleoporins or subcomplexes and will lead to the atomic resolution structures of individual human nucleoporins and/or their subcomplexes thus answering the various questions for the better understanding of nuclear pore complex assembly and functions. Given that the various Nups are involved in several diseases (Nofrini et al., 2016), better structural information will undoubtedly decipher the mechanisms and cure for such diseases.

## Materials and Methods-include

- Method details
  - Cloning and protein expression
  - Protein purification
  - SEC-MALS analysis
  - Western Blotting and Pull-down assay
  - Cryo-EM specimen preparation and data acquisition
  - Cryo-EM data analysis
  - Model building and refinement
  - Immunofluorescence analysis
- Contact for reagent and resource sharing
- Data and software availability

## Supplemental Information

Supplemental information includes seven figures and one table.

## Author Contributions

Conceptualization and supervision: R.C.; designing, execution and data analysis: S.N., J.S. and R.C.; Manuscript writing: S.N., J.S. and R.C. S.N. performed structural analysis and J.S. performed protein interaction analysis.

## Acknowledgments

We thank the ESRF CM01, and Dr. Eaazhisai Kandiah for assistance in EM data collection. We acknowledge the kind help of Dr. M. Kumar and Dr. M. Banerjee at IIT, Delhi in grid vitrification and Dr. Vinoth Kumar at the National Electron Cryo-Microscopy facility at the Bangalore Life Sciences Cluster (DBT/PR12422/MED/31/287/2014). We are thankful to A. Kembhavi and K. Waghmare, at IUCAA, Pune for helping with data transfer. This work was supported by the DBT grant (DBT/PR/26398) awarded to R.C and NCCS intramural funds. S.N. and J.S. thank UGC (22/12/2013(ii) EU-V/307720 and Dec-2018/808/354794, respectively) for the Ph.D. research fellowship. We thank members of the LSB group and K Saikrishnan (IISER Pune) for various suggestions and for critically reading the manuscript.

## Declaration of interests

The authors declare no competing interests.

## Supplementary Data

**Figure S1 (Related to Figures 1, 2, 3, 6, and 7):**
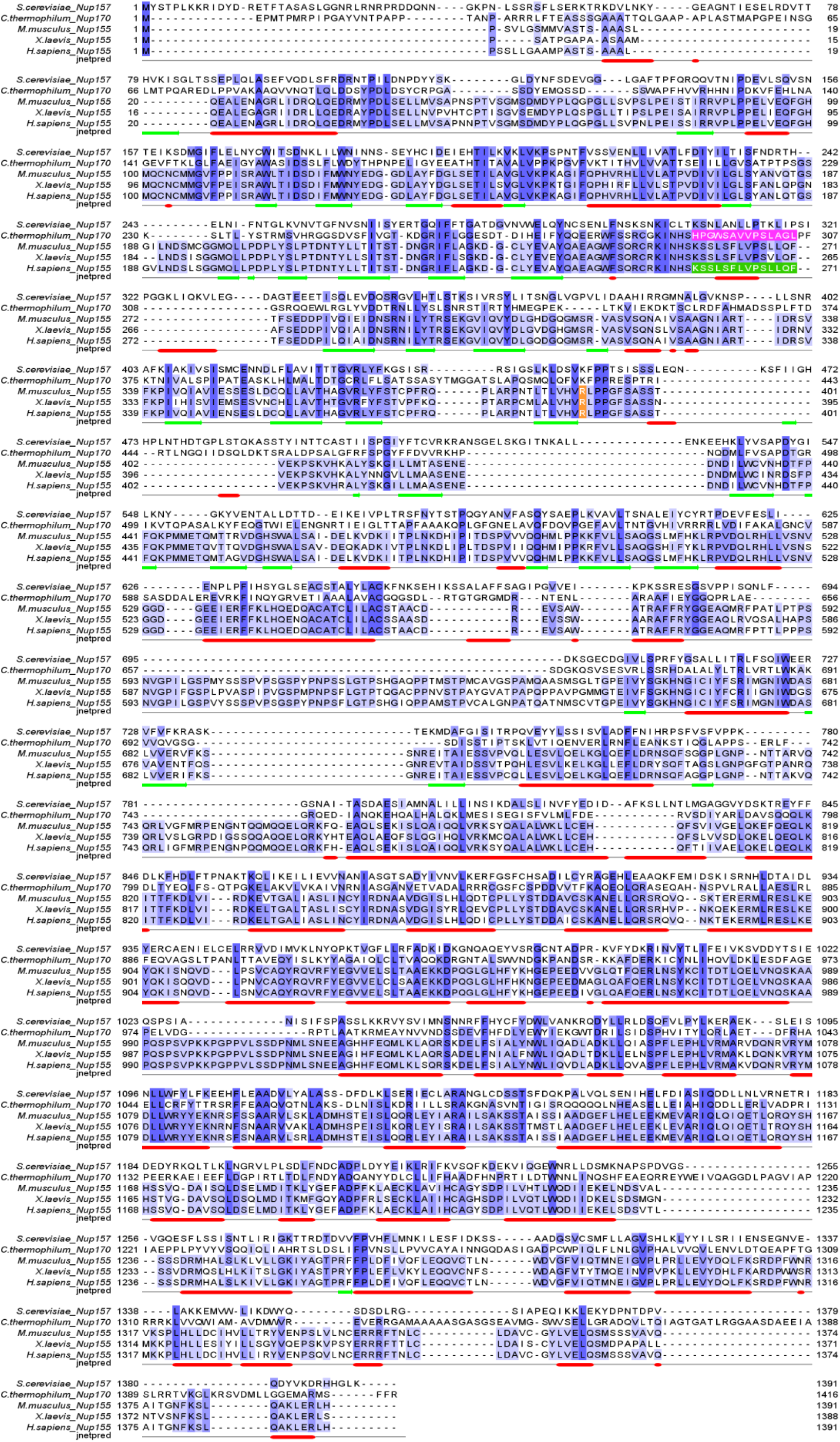
Structure-guided MSA of Nup155. Sequences from three vertebrate species (*Mus musculus, Xenopus laevis*, and *Homo sapiens*) are aligned with two lower eukaryotic species (*Saccharomyces cerevisiae* and *Chaetomium thermophilum)* using PROMALS 3D. Secondary structures are represented, α-helices as red bars, β-sheets as green arrow bars, and unstructured regions as black lines. The aligned residues are colored on the basis of sequence similarity from dark blue (100% identity), to mid-blue (50% similarity), to light blue (30% similarity), to white (below 30% similarity). The ALPS motif (membrane binding) is highlighted in green color on HsNup155 and pink color on CtNup170. The R391 location in human is highlighted with orange color.

**Figure S2 (Related to Figure 1):**
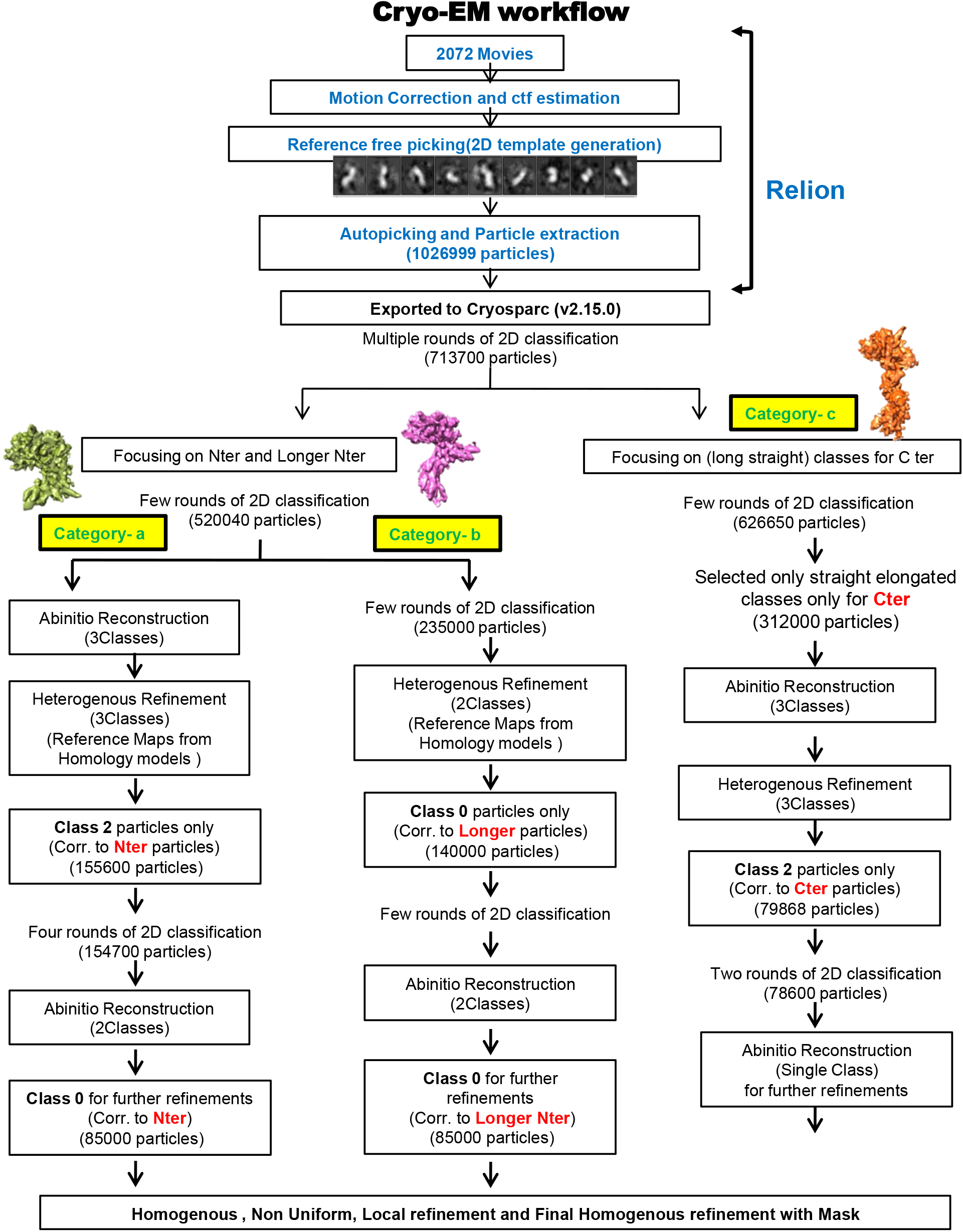
Overall Cryo-EM workflow followed to classify 3 different types of particles. Initially picked particles were cleaned and processed to pull out crescent-shaped particles (Category ‘a’) and question mark-shaped longer N-terminus (Category b) separately. Category c particles were filtered to obtain particles corresponding to elongated shaped C-terminus fragment.

**Figure S3 (Related to Figures 2,3 and 4):**
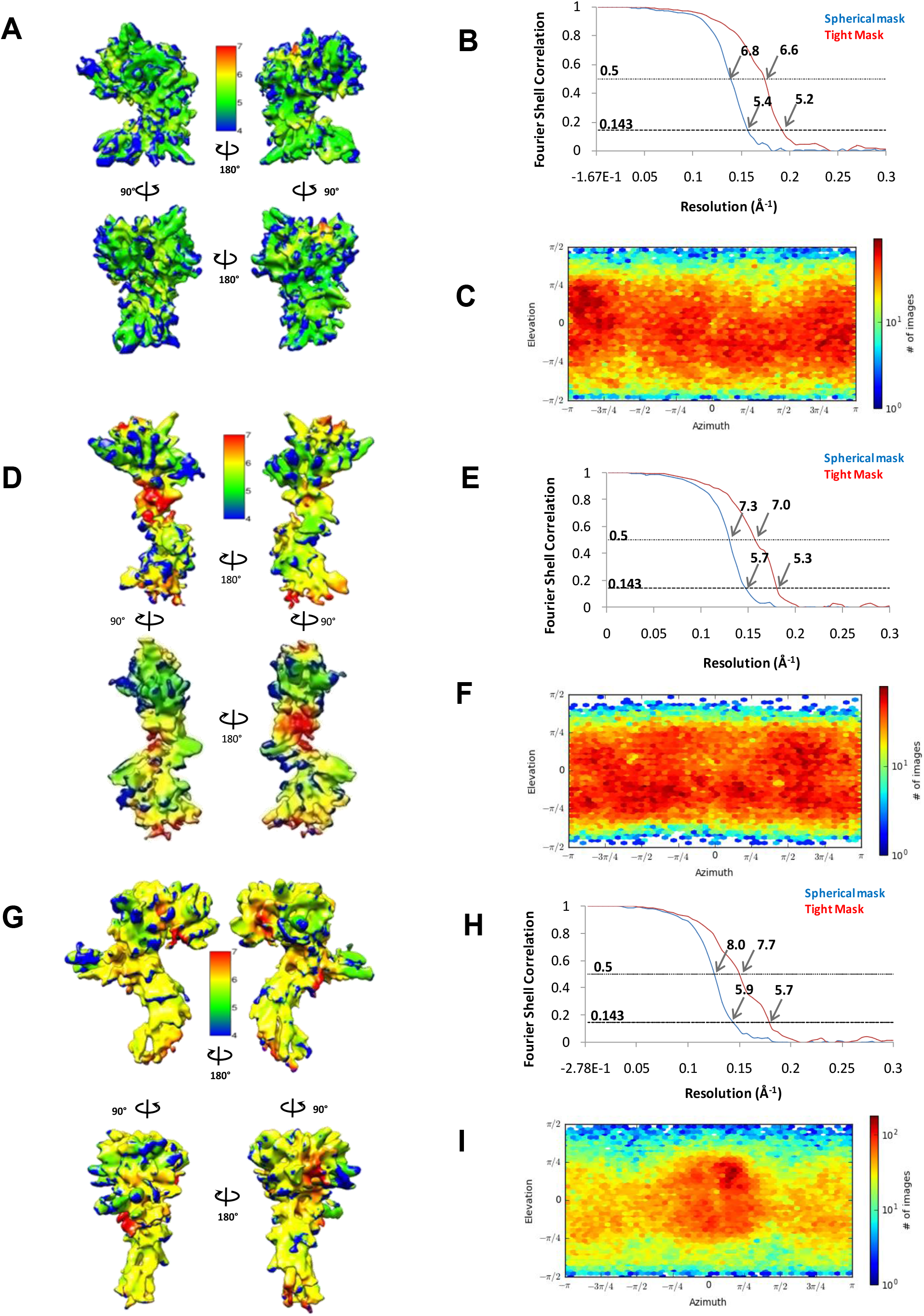
Cryo-EM analysis of *Hs*Nup155 structures. (A, D, and G) Shows the final density map for N-terminus, C-terminus, and longer N-terminus respectively, colored according to local resolution estimated by RESMAP. (B, E, and H) FSC curves for the density maps of Nup155 (19-863, 864-1337, and 19-1069 respectively) showing resolution as per Gold standard 0.143 FSC. (C, F and I) Viewing direction distribution of particles used for 3D reconstruction over azimuth and elevation angle for N-terminus, C-terminus, and longer N-terminus respectively.

**Figure S4 (Related to Figures 2 and 4):**
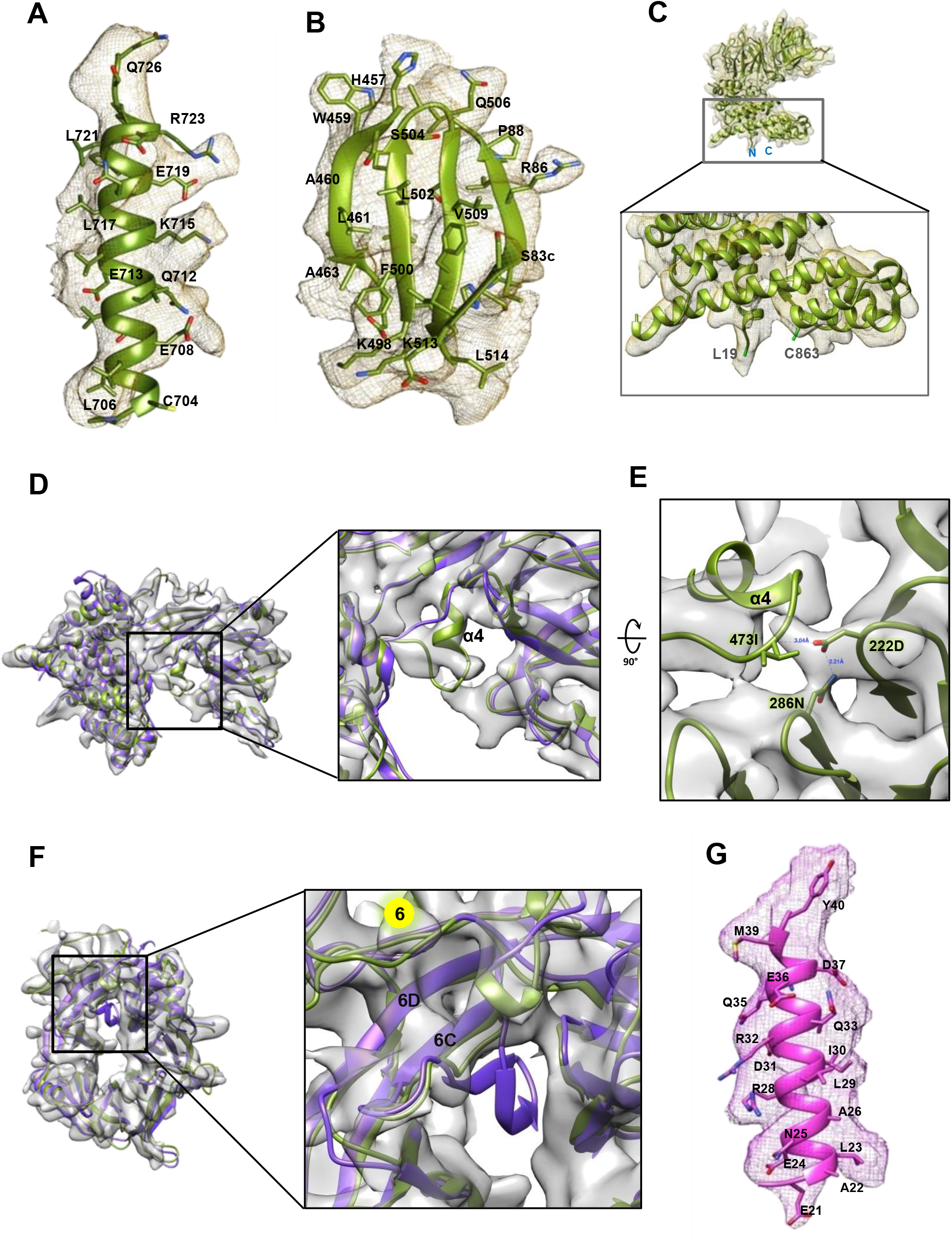
Cryo-EM analysis of *Hs*Nup155 Crescent-shaped domain. (A and B) CryoEM density maps showing good fitting for helix α11 and β-propeller 7^th^ blade respectively. (C) The density map showing termini residue boundaries from 19-863 residues. (D) Position of α4-helix, unique to *Hs*Nup155, at interaction interface of β-propeller and α-solenoid domain. *Hs*Nup155 is shown with olive green color superpositioned with *Ct*Nup170 structure (PDB ID:5HAX) in purple color. (E) Interaction of residue 473I at α4-helix with interstrand loops (with 222D and 286N) (F) Cryo-EM density along with 6^th^ blade of *Hs*Nup155 β-propeller domain (olive green color) having only 2 strands, superpositioned with *Ct*Nup170 structure in purple color with 4 strands in the corresponding blade. (G) CryoEM density map and fit for helix α1 showing overall fitting

**Figure S5 (Related to Figure 2):**
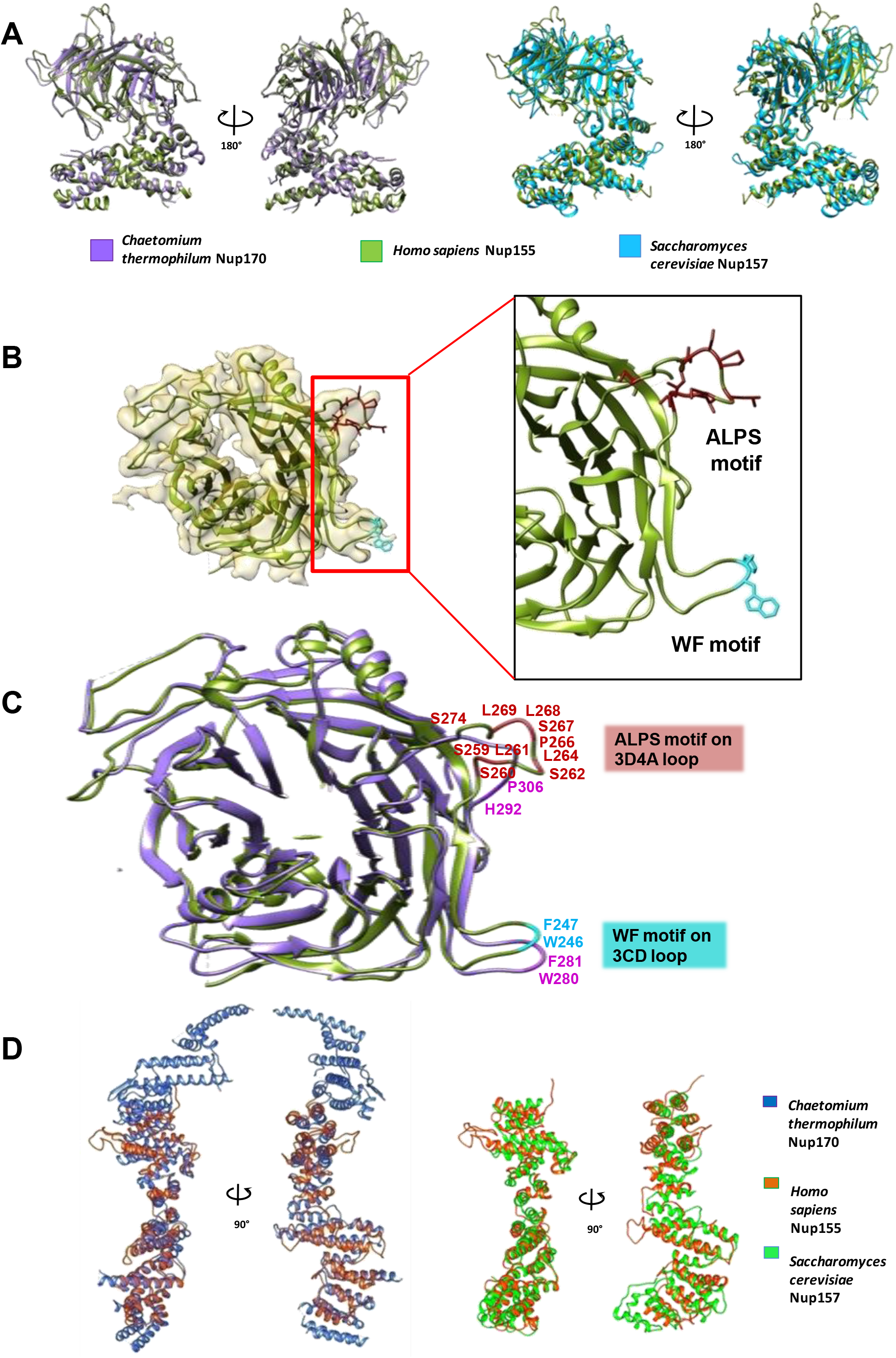
Comparison of *Hs*Nup155 N- and C-terminus domains with homologous structures. (A) Superposition of *Hs*Nup155 N-terminus (olive green color) Cryo-EM structure with Crystal structures of *Ct*Nup170 (PDB: 5HAX) in purple color and *Sc*Nup157 (PDB: 4MHC) in sky blue color with an RMSD (Cα) of 4.453 and 6.358 respectively. (B) Location of WF (cyan color) and ALPS motifs (dark red color) on interstrand and interblade loops respectively. (C) Superposition of *Hs*Nup155 β-propeller domain on 5HAX (purple color). The WF motif on 3CD loop is well conserved in *Ct* and *Hs*. The ALPS motif from *Ct* has homologous loop on 3D4A loop, rich in serine, lysine and proline in *Hs*Nup155. (D) Superposition of *Hs*Nup155 C-terminus (orange color) Cryo-EM structure with Crystal structures of *Ct*Nup170 (PDB: 5HB1) in blue color and *Sc*Nup170 (PDB: 3I5P) in light green color with an RMSD (Cα) of 4.793 and 6.846 respectively.

**Figure S6 (Related to Figures 6 and 7):**
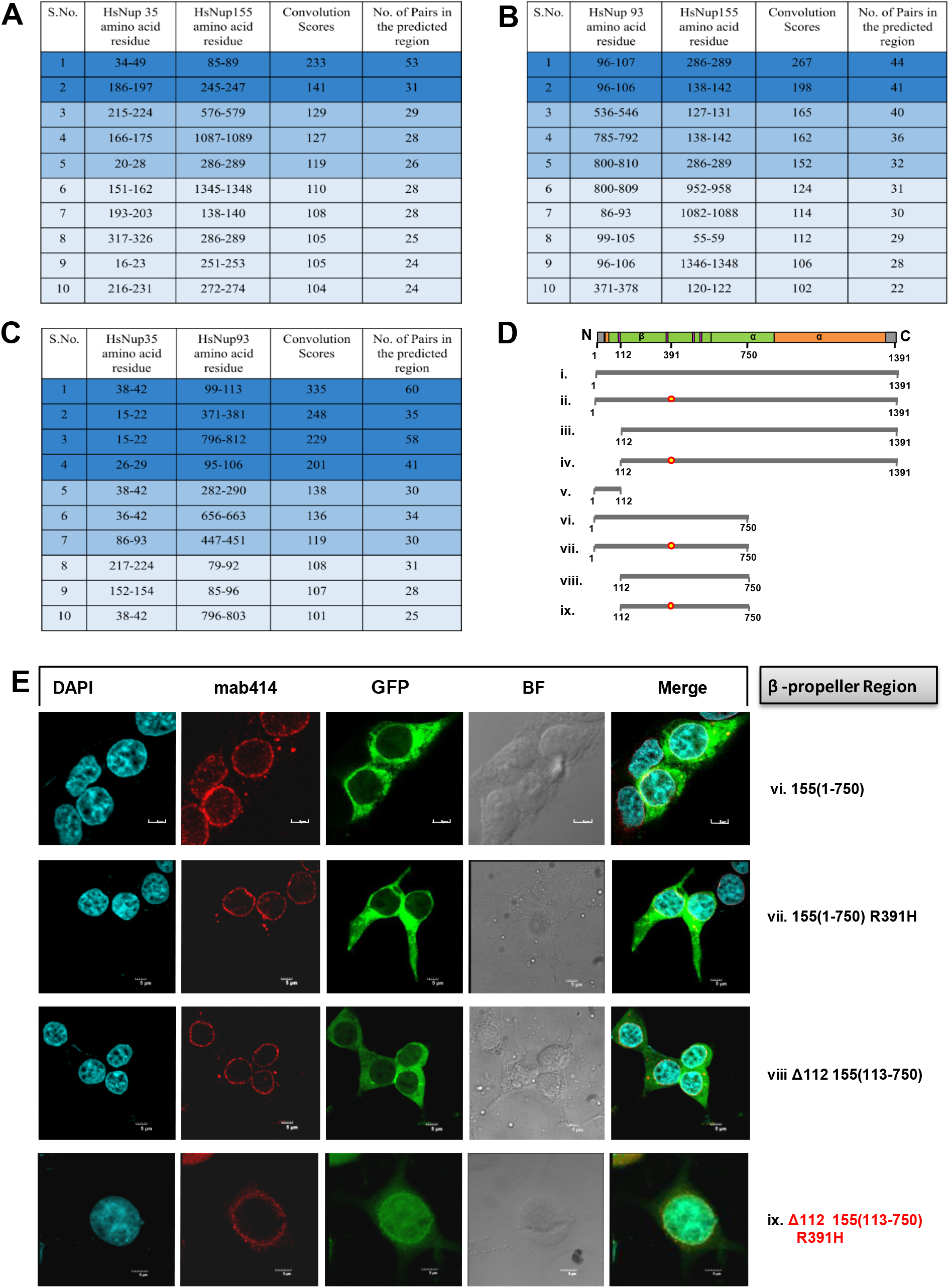
Prediction of Interface regions for *Hs*Nup93-*Hs*Nup155-*Hs*Nup35 using CoRNeA. (A-C) Tables summarizing top 10% interacting residue pairs with the convolution scores obtained from CoRNeA for Nup35•Nup155 (A), Nup93•Nup155 (B) and Nup35•Nup93 pairs (C) interactions respectively. The top two rows represents highest scoring residue pair (Dark blue), rows 3-5 intermediate scoring pairs (sky blue) and rows 6-10 lower scoring residue pairs (Teal blue). (D)Schematic representation of various deletion constructs used for confocal microscopy. The red circle over the line represents R391H mutation. (E) GFP fluorescence-based localization assays for b-propeller domain (1-750) of *Hs*Nup155; showing:vi: Nup155(1-750), vii:Nup155(1-750)R391H and viii:hsNup155(113-750) is localizing on NPC. However, ix: Nup155(112-750)R391H shows a mislocalization pattern.

**Figure S7 (Related to Figures 6 and 7):**
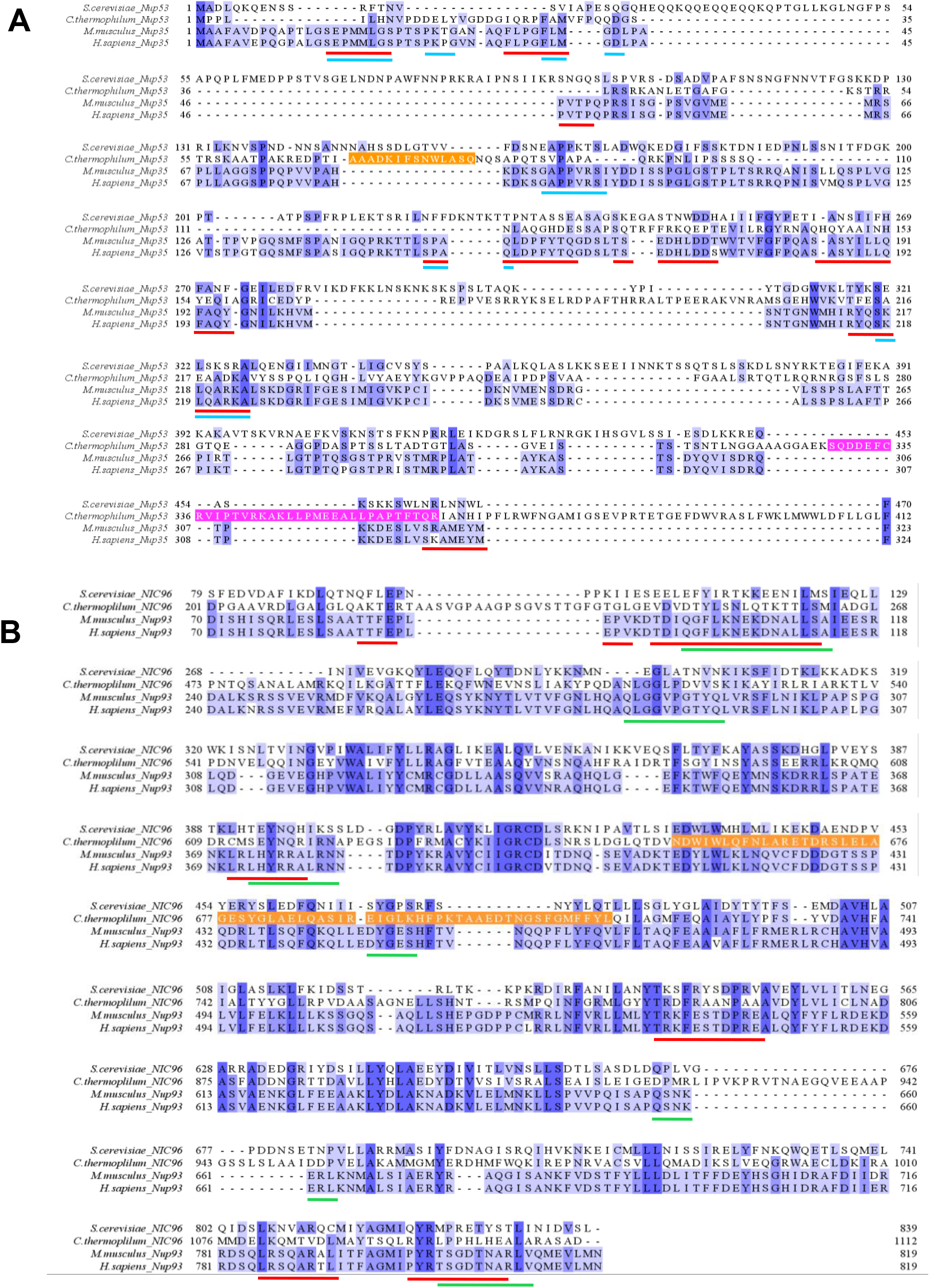
Interaction mapping on multiple sequence alignment of Nup35 (A) and Nup93 (B). Sequences from two vertebrate species (*Mus musculus*, and *Homo sapiens*) are aligned with two lower eukaryotic species (*Saccharomyces cerevisae* and *Chaetomium thermophilum)* using PROMALS 3D. The aligned residues are colored on the basis of sequence similarity from dark blue (100% identity), to mid-blue (50% similarity), to light blue (30% similarity), to white (below 30% similarity). (A) Nup35 alignment showing CoRNeA predicted interaction regions on *Hs*Nup35 are highlighted with colored bars below the sequence, Red color for Nup155 and cyan color for Nup93 interacting regions. Region in *Ct*Nup53 shown to interact with Nic96 and Nup170 in orange and pink color, respectively. (B) Nup93 alignment showing CoRNeA predicted interaction regions on *Hs*Nup93 are shown as colored bars; red color for Nup155 and green color for Nup35 interacting regions. Previously known *Ct*Nic96 is highlighted for the Nup53 interacting region in orange color.

## Materials and Methods

### Method details

#### Cloning and protein expression

Human NUP155 gene (UniProt accession no. 075694), full length (1391 residues) as well as different C-terminus and N-terminus truncated constructs were cloned in pEGFP-C1 vector (clontech), based on its secondary structure prediction by Psipred (McGuffin et al., 2000).

Suspension cultures of human embryonic kidney (HEK) 293F cells (purchased from Invitrogen) were maintained using 293 freestyle media (gibco) at 37°C with 8% CO_2_ and 80% humidity at 110rpm. The cells at 1.5*10^6^ confluence were transiently transfected with 1: 3 ratio of desired DNA and polyethylenimine (PEI) (Polysciences) complex and incubated at 37°C with 8% CO_2_, 80% humidity in shaking condition at 110rpm for 65-70 hours post-transfection.

#### Protein purification

For Purifying nanobody, the GST-tagged anti-GFP nanobody construct in pGEX-6P-1 vector (Addgene ID # 61838) was transformed to BL21 RIL (DE3) cells. The cells were cultured in LB containing 1% glucose and 1mM MgCl_2_ at 37°C until the OD reached 0.5. The culture was then induced with 0.5mM IPTG at 18°C for 12 hours. Harvested cells were lysed by sonication in a buffer containing 50mM Tris pH 8, 300mM NaCl, 1mM βMe, and 1mM PMSF. The cell lysate was centrifuged at 12,000 rpm for 45min and the supernatant was kept for binding with glutathione resin for 3hours at 4°C. The resin was washed with buffer containing 10mM Tris-Cl pH 8, 250mM NaCl, 1mM β-Me and then eluted with 10mM Tris-Cl pH 8, 250mM NaCl, 1mM β-Me and 10mM reduced glutathione. The elution fractions were further purified with size exclusion chromatography (SEC) using Superdex 200 16/600 pg column (GE Healthcare) in buffer containing 10mM Tris-Cl pH 8, 150mM NaCl, 1mM DTT and 0.5mM EDTA. The purified GST-tagged anti-GFP nanobody was then snap-frozen in liquid nitrogen and stored at -80°C. The purified GST-tagged anti-GFP nanobody allowed to bind to glutathione resin (Pierce) for 2 hours at 4°C in rocking condition followed by removing the unbound nanobodies by washing of the resin with a buffer containing 50mM Tris, 300mM NaCl, 1mM DTT, 0.5mM EDTA and 5% glycerol. This nanobody-bound resin was used to bind *Hs*Nup155 from cell lysate.

For purifying the N-terminus GFP-tagged *Hs*Nup155 pEGFP-C1 vector was used. HEK293F Cells over-expressed with *Hs*Nup155 were lysed by sonication in a buffer containing 50mM Tris pH 7.5, 300mM NaCl, 1%DDM (n-Dodecyl-β-D-maltoside) (Inalco), 1mM DTT, 0.5mM EDTA, 5% glycerol and protease inhibitor cocktail (Roche). The cell debris was removed from the lysate by centrifugation at 16,000 rpm for 30min. The supernatant was collected and kept for binding for 4 hours at 4°C with glutathione resin which was pre-bound with GST-tagged anti-GFP nanobodies. The resin was washed with 50mM Tris pH 7.5, 300mM NaCl, 0.1mM DDM, 1mM DTT, 0.5mM EDTA and 3% glycerol for 20 column volumes of the resin and then *Hs*Nup155 and anti-GFP nanobody complex was eluted with a buffer containing 50mM Tris pH 7.5, 300mM NaCl, 0.05mM DDM, 1mM DTT, 1mM EDTA, 1% glycerol and 10mM reduced glutathione.

The GST tag present in *Hs*Nup155 and anti-GFP nanobody complex was removed by digesting the protein with PreScission protease for 12hours at 4°C in rocking condition. The N-terminus GFP-tagged HsNup155 bound with anti-GFP nanobody was further purified by size-exclusion chromatography in 50mM Tris pH 7.5, 150mM NaCl, 0.05mM DDM, 1mM DTT, 1mM EDTA and 1% glycerol.

#### SEC-MALS analysis of the purified sample

The purified protein was analyzed by Size Exclusion Chromatography (SEC) (Superdex 200 10/30 GL column; GE Healthcare) coupled with multi-angle light scattering (MALS) instrument (Wyatt Technology). The Wyatt Dawn Heleos-II light scattering detector and Wyatt Optilab T-rex refractive index detector were connected in-line with the chromatography system for simultaneous MALS measurements. 0.05mg of purified protein was injected with a buffer containing 50mM Tris pH 7.5, 150mM NaCl, 0.05mM DDM, 1mM DTT, 1mM EDTA and 1% glycerol. The BSA was used as standard (2 mg/mL; Pierce) and data was analyzed with Astra software (Wyatt Technologies). All experiments were performed at room temperature.

#### Western Blotting

The protein sample from the same batch which was used for Cryo-EM grid preparation was loaded on SDS-PAGE gel and separated by electrophoresis. The protein bands were blotted onto PVDF membrane and probed with anti-Nup155 antibody (abcam EPR17111), which is specific for C-terminus residues (900-1050 aa). The duplicate blot was separately probed with anti-GFP antibody (Sigma G1546), to detect the GFP at the N-terminus of *Hs*Nup155. The blots were further probed with Secondary anti-rabbit (Promega W401B) and secondary anti-mouse (Sigma A3673) antibodies respectively.

#### Pull down assay

**Pulldown assay**-To identify the interacting regions of the *Hs*Nup93-*Hs*Nup35 Complex,GST based pulldown method was used. *Hs*Nup93 (96-819) and HsNup93 (1-150) in pGEX4T1 vector was co-transformed with *Hs*Nup35 (1-305) in pET28a vector respectively. Similarly, *Hs*Nup93 (96-819)in pGEX4T1 vector was co-transformed with*Hs*Nup155(1-515)in pET28a vector in *E*.*coli* BL21 (DE3) RIL cells. The transformed cells were cultured in 1 liter of LB broth containing corresponding antibiotics at 37°C and induced with 0.5mM IPTG at OD_600_ about 0.4-0.6 and incubated at 18°C post-induction for 12-14 hrs. Harvested cells were lysed in lysis buffer (20mM HEPES pH7.4,200mM NaCl, 0.5mM EDTA, 1mMDTT, 1mM PMSF, 5mM β-Me and 2% Glycerol)Lysates were incubated with 500ul of GSH-Sepharose (Pierce) beads for 3hrs with rotation at 4°C. Beads were washed with 40 CV of Wash Buffer (20mM HEPES, 100mM NaCl,0.5mM EDTA). Complexes were eluted with 1 CV of Elution Buffer (20mM HEPES, 150mM NaCl, 0.5mM EDTA,10mM Reduced L-Glutathione,1mM PMSF,1% Glycerol).The pulled-out complexes were resolved on denaturing SDS PAGE. Western Blot analysis was carried out using Anti-GST(1:3000) antibody (Sigma #G1160) and Anti-His(1:3000) antibody(Sigma #H1029).HRP conjugated mouse IgG was used at 1:4000 dilution(Sigma #A3673) for developing the signal. Images were recorded using Imager800 (GE Healthcare).

#### Cryo-EM specimen preparation and data acquisition

Specimens were prepared of vitrified N-GFP *Hs*Nup155 and anti-GFP nanobody complex protein sample by placing 3µl of protein at a concentration of 1.3mg/ml sample on a freshly glow-discharged Quantifoil holey carbon grid (R 2/1, 200-mesh ultra foil Au grid). After 3sec of blotting with zero blot force in 100% humidity at 18°C, the grid was vitrified by plunging into liquid ethane using a Vitrobot (FEI; Mark IV). Storage and transport of the prepared specimen were done in liquid nitrogen.

Cryo-EM data collection was carried out on Titan Krios microscope operated at 300KV and equipped with a K2 detector with a post-column energy filter. Micrographs were recorded in super-resolution mode at a magnified nominal pixel size of 0.823 Å and defocus ranging from −1.0 to −2.5 μm with a step size of 0.2 μm. A total of 7300 movies were collected in counting mode with a dose rate of 1.15 e^-^ per Å^2^, 50 dose-fractionated frames were collected which counts for a total dose of 57.6 e^-^ per Å (Kandiah et al., 2019).

#### Cryo-EM data analysis

The movies were aligned, and dose-weighted to correct for movement during imaging and account for radiation damage via Motioncor2 (Zheng et al., 2017). The CTF parameters for each micrograph were determined by Gctf (Zhang, 2016). A total of 2,072 movies were selected manually after excluding the micrographs with ice contamination and drifting issues. Nearly 5000 particles were picked manually and 2D classified to generate templates for automated particle picking in RELION 3.1 beta (Scheres, 2012). A total of ∼10,27,000 particles were autopicked and exported to cryoSPARC V2 (Punjani et al., 2017) for further processing (Figure S2). Particles were subjected to several rounds of reference-free 2D classification followed by manual inspection and selection of classes with some visible features. This process yielded a stack of around 7,13,700 cleaned particles that were subjected to further 2D classification and ab initio 3D reconstruction separately thrice to scrutinize 3 different types of particles present over the grid (Figure 1E-G).

As per the western blot analysis, three bands corresponding to different types of particles of *Hs*Nup155 were present on the grid: Longer N-terminus (Maybe Full length (1-1391)), N-terminus and C-terminus. The HsNup155 full-length protein got cleaved into N-terminus (1-863) and C-terminus (864-1391). The boundaries of the residues for cleavage were found after solving the cryo-EM data and fitting the model into N-terminus and C-terminus EM density maps. Based on the shapes of these classes of particles, the initially cleaned 713700 particles were processed repeatedly for the scrutiny of particles corresponding to all 3 types of particles present on the grid. The N-terminus particles were smaller head regions of question mark-shaped particles corresponding to longer N-terminus particles were further cleaned to 520040 particles focusing on N-terminus and longer N-terminus classified as “Category a and b respectively”. While the particles corresponding to C-terminus were more elongated and straight: “Category c” particles.

##### Separation of particles corresponding to 3 different classes

**cleaned 520040** particles were processed twice focusing on **N-terminus “Category a”** and **Longer N-terminus “Category b”** separately.

1. For obtaining **N-terminus** EM density, 520040 particles were used for ab initio-reconstruction into 3classes. The EM density maps obtained were similar to 3 types of classes corresponding to N-terminus, longer N-terminus and C-terminus. To further scrutinize the particles specific to N-terminus only, heterogenous refinement was performed into 3 classes in which the homogenous refined volume maps from ab initio reconstruction were used while the N-terminus corresponding volume map (crescent-shaped) was generated on *Hs*Nup155 N-terminus model on PDB: 5HAX as a template. The 155600 particles obtained in class 2 were further used for 2D classification followed with ab initio reconstruction into 2 classes. Class 0 having ∼85000 particles corresponding to N-terminus was further refined.
2. For obtaining **Longer N-terminus** EM density, 520040 particles were 2D classified and cleaned to segregate longer question mark-shaped particles. Final 235000 particles were further subjected to heterogeneous refinement into 2 classes using volume maps built on homology model for longer N-terminus (using PDBs 5HAX and 5HB1 as template) for class 1 and to separate elongated C-terminus corresponding particles, volume map on homology model for C-terminus (on PDB 5HB1 as template) for class 2 was used. Class 0 having 130600 particles was further used for 2D classification and ab initio reconstruction into 2 classes. Class 0 having ∼85000 particles corresponding to longer N-terminus was further refined. The initial 713700 particles were again 2D classified and cleaned to 626650 particles and kept in **“Category c”** for further extraction of C-terminus corresponding particles from it.
3. For obtaining **C-terminus** EM density, 626650 particles were 2D classified and only straight elongated 2D classes were subset classified containing 312000 particles. These particles were further subjected to ab initio reconstruction into 3 classes followed by heterogeneous refinement into 3 classes using homogenous refined maps from ab initio reconstruction. 79868 particles obtained in class 2 corresponding to straight C-terminus were 2D classified and cleaned to 78600 particles followed with ab initio reconstruction in a single class which was further refined.

For all the 3 classes, EM maps obtained were used for Homogenous 3D refinement in C1 symmetry, followed with the non-uniform and local refinement with a specific mask on homology model and a final homogenous refinement with the specific mask as implemented in CryoSPARC V2 workflow, yielded final maps to a resolution of ∼ 5.2 Å ∼ 5.7 Å and ∼5.3 Å (GSFSC_0.143_) for N-terminus, Longer N-terminus and C-terminus respectively.

#### Model building and refinement

Each of the N-terminus and C-terminus domains modeled via Swiss-model (Schwede, 2003) using PDB: 5HAX and 5HB1 for N and C-terminus respectively. The final model was built by the rigid-body fitting of domains from homology models into the EM map in COOT. A model for full length (1-1391) was generated by combining the solved structures of N-terminus and C-terminus following the overlapping density obtained via EM density map of longer N-terminus. The fits were improved by multiple rounds of real-space refinement in Phenix (Adams et al., 2010; Afonine et al., 2018). The CC between models and EM map of N-terminus, longer N-terminus and C-terminus were 0.56, 0.6 and 0.54 respectively, indicative of a reasonable fit at the present resolution. The final model has good stereochemistry, as evaluated using MolProbity (Davis et al., 2004) (Table 1). All of the figures were prepared with UCSF Chimera (Pettersen et al., 2004).

#### Immunofluorescence analysis of HsNup155 localization

HEK293 cells grown in DMEM media (Gibco) supplied with 10%FBS and 2mM L-Glutamine were seeded onto coverslips in 24 well plates and allowed to adhere and grow overnight at 37°C incubator supplied with 5% CO_2_. The cells were transfected with 1: 3 ratio of desired DNA and polyethylenimine (PEI) (Polysciences) complex and incubated the cells for 35-36 hours at 37°C before preparation of slides for confocal microscopy. The cells on the coverslips were washed with DPBS (Gibco) and fixed with 4% paraformaldehyde for 20 min. Cells were again washed with DPBS and incubated with 0.1% glycine in DPBS for 5 min. 0.1% triton x-100 (sigma) in DPBS was used for permeabilization followed by washing thrice with DPBS. 10% FBS was used for blocking at 4°C overnight. Coverslips were incubated overnight with primary antibody mAb414 (1:300; Abcam #ab24609) for 10-12hrs at 4°C. Samples were washed thrice with DPBS before incubating it with secondary antibody Alexa Fluor 594 (1:1000, ThermoFisher #A21203) for 1hour at RT. Samples were washed with DPBS and stained with DAPI for 10min at RT. The coverslips were finally washed with DPBS and mounted on glass slides using Vectasheild antifade mounting medium (Vector Laboratories) to maintain samples until imaging. Images were acquired on a confocal Olympus FV3000 microscope.

## Contact for reagent and resource sharing

Further information and requests for resources and reagents should be directed to and will be fulfilled by the Lead Contact Radha Chauhan.

## Data and software availability

Cryo-EM maps of the *Hs*Nup155 N-terminus, C-terminus and longer N-terminus with its associated atomic models are deposited in the wwPDB OneDep System under EMD accession codes EMD-31382, 31383 and 31384 respectively and PDB ID codes-7EYE, 7EYF and 7EYQ respectively.

## Notes

### Competing Interest Statement

The authors have declared no competing interest.

## References

1. Afanasieva, E., Chaudhuri, I., Martin, J., Hertle, E., Ursinus, A., Alva, V., Hartmann, M.D., and Lupas, A.N. (2019). Structural diversity of oligomeric β-propellers with different numbers of identical blades. Elife 8.

2. Alber, F., Dokudovskaya, S., Veenhoff, L.M., Zhang, W., Kipper, J., Devos, D., Suprapto, A., Karni-Schmidt, O., Williams, R., Chait, B.T., et al. (2007). The molecular architecture of the nuclear pore complex. Nature 450.

3. Amlacher, S., Sarges, P., Flemming, D., Van Noort, V., Kunze, R., Devos, D.P., Arumugam, M., Bork, P., and Hurt, E. (2011). Insight into structure and assembly of the nuclear pore complex by utilizing the genome of a eukaryotic thermophile. Cell 146, 277–289.

4. Andersen, J.H., Andreasen, L., and Olesen, M.S. (2020). Atrial fibrillation—a complex polygenetic disease. Eur. J. Hum. Genet.

5. Anter, E., Jessup, M., and Callans, D.J. (2009). Atrial Fibrillation and Heart Failure. Circulation 119.

6. Von Appen, A., Kosinski, J., Sparks, L., Ori, A., DiGuilio, A.L., Vollmer, B., Mackmull, M.T., Banterle, N., Parca, L., Kastritis, P., et al. (2015). In situ structural analysis of the human nuclear pore complex. Nature 526, 140–143.

7. Beck, M. (2004). Nuclear Pore Complex Structure and Dynamics Revealed by Cryoelectron Tomography. Science (80-.). 306.

8. Beck, M., and Hurt, E. (2017). The nuclear pore complex: Understanding its function through structural insight. Nat. Rev. Mol. Cell Biol. 18, 73–89.

9. Beck, M., Lučić, V., Förster, F., Baumeister, W., and Medalia, O. (2007). Snapshots of nuclear pore complexes in action captured by cryo-electron tomography. Nature 449.

10. Belgareh, N., Rabut, G., Baï, S.W., van Overbeek, M., Beaudouin, J., Daigle, N., Zatsepina, O. V., Pasteau, F., Labas, V., Fromont-Racine, M., et al. (2001). An evolutionarily conserved NPC subcomplex, which redistributes in part to kinetochores in mammalian cells. J. Cell Biol. 154.

11. Breuer, M., and Ohkura, H. (2015). A negative loop within the nuclear pore complex controls global chromatin organization.

12. Bui, K.H., Von Appen, A., Diguilio, A.L., Ori, A., Sparks, L., Mackmull, M.T., Bock, T., Hagen, W., Andrés-Pons, A., Glavy, J.S., et al. (2013). Integrated structural analysis of the human nuclear pore complex scaffold. Cell 155, 1233–1243.

13. Busayavalasa, K., Chen, X., Farrants, A.K.Ö., Wagner, N., and Sabri, N. (2012). The Nup155-mediated organisation of inner nuclear membrane proteins is independent of Nup155 anchoring to the metazoan nuclear pore complex. J. Cell Sci. 125, 4214–4218.

14. Casañal, A., Shakeel, S., and Passmore, L.A. (2019). Interpretation of medium resolution cryoEM maps of multi-protein complexes. Curr. Opin. Struct. Biol. 58.

15. Chen, C.K.-M., Chan, N.-L., and Wang, A.H.-J. (2011). The many blades of the β-propeller proteins: conserved but versatile. Trends Biochem. Sci. 36.

16. Chopra, K., Bawaria, S., and Chauhan, R. (2019). Evolutionary divergence of the nuclear pore complex from fungi to metazoans. Protein Sci. 28.

17. Chopra, K., Burdak, B., Sharma, K., Kembhavi, A., Mande, S.C., and Chauhan, R. (2020). CoRNeA: A Pipeline to Decrypt the Inter-Protein Interfaces from Amino Acid Sequence Information. Biomolecules 10.

18. Cronshaw, J.M., Krutchinsky, A.N., Zhang, W., Chait, B.T., and Matunis, M.L.J. (2002). Proteomic analysis of the mammalian nuclear pore complex. J. Cell Biol. 158, 915–927.

19. Eibauer, M., Pellanda, M., Turgay, Y., Dubrovsky, A., Wild, A., and Medalia, O. (2015). Structure and gating of the nuclear pore complex. Nat. Commun. 6.

20. Eisenhardt, N., Redolfi, J., and Antonin, W. (2014). Interaction of Nup53 with Ndc1 and Nup155 is required for nuclear pore complex assembly. J. Cell Sci. 127.

21. Fahrenkrog, B., and Aebi, U. (2003). The nuclear pore complex: nucleocytoplasmic transport and beyond. Nat. Rev. Mol. Cell Biol. 4.

22. Franz, C., Askjaer, P., Antonin, W., Iglesias, C.L., Haselmann, U., Schelder, M., De Marco, A., Wilm, M., Antony, C., and Mattaj, I.W. (2005). Nup155 regulates nuclear envelope and nuclear pore complex formation in nematodes and vertebrates. EMBO J. 24, 3519–3531.

23. Frey, S., Richter, R.P., and Görlich, D. (2006). FG-rich repeats of nuclear pore proteins form a three-dimensional meshwork with hydrogel-like properties. Science (80-.). 314, 815–817.

24. Galy, V., Mattaj, I.W., and Askjaer, P. (2003). Caenorhabditis elegans Nucleoporins Nup93 and Nup205 Determine the Limit of Nuclear Pore Complex Size Exclusion In Vivo. Mol. Biol. Cell 14.

25. Grandi, P., Doye, V., and Hurt, E.C. (1993). Purification of NSP1 reveals complex formation with “GLFG” nucleoporins and a novel nuclear pore protein NIC96.

26. Grandi, P., Schlaich, N., Tekotte, H., and Hurt1, E.C. (1995). Functional interaction of Nic96p with a core nucleoporin complex consisting of Nsplp, Nup49p and a novel protein Nup57p.

27. Guan, T., Müller, S., Klier, G., Panté, N., Blevitt, J.M., Haner, M., Paschal, B., Aebi, U., and Gerace, L. (1995). Structural analysis of the p62 complex, an assembly of O-linked glycoproteins that localizes near the central gated channel of the nuclear pore complex. Mol. Biol. Cell 6.

28. Han, M., Zhao, M., Cheng, C., Huang, Y., Han, S., Li, W., Tu, X., Luo, X., Yu, X., Liu, Y., et al. (2019). Lamin A mutation impairs interaction with nucleoporin NUP155 and disrupts nucleocytoplasmic transport in atrial fibrillation. Hum. Mutat. 40, 310–325.

29. Hawryluk-Gara, L.A., Shibuya, E.K., and Wozniak, R.W. (2005). Vertebrate Nup53 Interacts with the Nuclear Lamina and Is Required for the Assembly of a Nup93-containing Complex ? D. Mol. Biol. Cell 16, 2382–2394.

30. Hawryluk-Gara, L.A., Platani, M., Santarella, R., Wozniak, R.W., and Mattaj, I.W. (2008). Nup53 Is Required for Nuclear Envelope and Nuclear Pore Complex Assembly. Mol. Biol. Cell 19.

31. Hu, T., Guan, T., and Gerace, L. (1996). Molecular and functional characterization of the p62 complex, an assembly of nuclear pore complex glycoproteins. J. Cell Biol. 134.

32. Khan, A.U., Qu, R., Ouyang, J., and Dai, J. (2020). Role of Nucleoporins and Transport Receptors in Cell Differentiation. Front. Physiol. 11.

33. Kopec, K.O., and Lupas, A.N. (2013). β-Propeller Blades as Ancestral Peptides in Protein Evolution. PLoS One 8.

34. Kosinski, J., Mosalaganti, S., Von Appen, A., Teimer, R., Diguilio, A.L., Wan, W., Bui, K.H., Hagen, W.J.H., Briggs, J.A.G., Glavy, J.S., et al. (2016). Molecular architecture of the inner ring scaffold of the human nuclear pore complex. Science (80-.). 352, 363–365.

35. Leonard, R.J., Preston, C.C., Gucwa, M.E., Afeworki, Y., Selya, A.S., and Faustino, R.S. (2020). Protein Subdomain Enrichment of NUP155 Variants Identify a Novel Predicted Pathogenic Hotspot. Front. Cardiovasc. Med. 7.

36. Lin, D.H., and Hoelz, A. (2019). The Structure of the Nuclear Pore Complex (An Update). Annu. Rev. Biochem. 88.

37. Lin, D.H., Stuwe, T., Schilbach, S., Rundlet, E.J., Perriches, T., Mobbs, G., Fan, Y., Thierbach, K., Huber, F.M., Collins, L.N., et al. (2016). Architecture of the symmetric core of the nuclear pore. Science (80-.). 352.

38. Loïodice, I., Alves, A., Rabut, G., van Overbeek, M., Ellenberg, J., Sibarita, J.-B., and Doye, V. (2004). The Entire Nup107-160 Complex, Including Three New Members, Is Targeted as One Entity to Kinetochores in Mitosis. Mol. Biol. Cell 15.

39. Maimon, T., Elad, N., Dahan, I., and Medalia, O. (2012). The human nuclear pore complex as revealed by cryo-electron tomography. Structure 20, 998–1006.

40. Marzlin, K.M. (2020). Atrial Fibrillation and Heart Failure. AACN Adv. Crit. Care 31.

41. Mio, K., and Sato, C. (2018). Lipid environment of membrane proteins in cryo-EM based structural analysis. Biophys. Rev. 10, 307–316.

42. Mitchell, J.M., Mansfeld, J., Capitanio, J., Kutay, U., and Wozniak, R.W. (2010). Pom121 links two essential subcomplexes of the nuclear pore complex core to the membrane. J. Cell Biol. 191, 505–521.

43. Mosalaganti, S., Kosinski, J., Albert, S., Schaffer, M., Strenkert, D., Salomé, P.A., Merchant, S.S., Plitzko, J.M., Baumeister, W., Engel, B.D., et al. (2018). In situ architecture of the algal nuclear pore complex. Nat. Commun. 9.

44. Murzin, A.G. (1992). Structural principles for the propeller assembly of β-sheets: The preference for seven-fold symmetry. Proteins Struct. Funct. Genet. 14.

45. Nakano, H., Wang, W., Hashizume, C., Funasaka, T., Sato, H., and Wong, R.W. (2011). Unexpected role of nucleoporins in coordination of cell cycle progression. Cell Cycle 10.

46. Nanni, S., Re, A., Ripoli, C., Gowran, A., Nigro, P., D’Amario, D., Amodeo, A., Crea, F., Grassi, C., Pontecorvi, A., et al. (2016). The nuclear pore protein Nup153 associates with chromatin and regulates cardiac gene expression in dystrophic mdx hearts. Cardiovasc. Res. 112, 555–567.

47. Nofrini, V., DI Giacomo, D., and Mecucci, C. (2016). Nucleoporin genes in human diseases. Eur. J. Hum. Genet. 24, 1388–1395.

48. Nudelman, I., Kim, S.J., Fernandez-Martinez, J., Shi, Y., Zhang, W., Raveh, B., Herricks, T., Slaughter, B.D., Hogan, J., Upla, P., et al. (2018). Integrative Structure and Functional Anatomy of a Nuclear Pore Complex. Microsc. Microanal. 24, 1212–1213.

49. Ori, A., Banterle, N., Iskar, M., Andrés-Pons, A., Escher, C., Khanh Bui, H., Sparks, L., Solis-Mezarino, V., Rinner, O., Bork, P., et al. (2013). Cell type-specific nuclear pores: A case in point for context-dependent stoichiometry of molecular machines. Mol. Syst. Biol. 9.

50. Parvez, B., and Darbar, D. (2011a). The “missing” link in atrial fibrillation heritability. J. Electrocardiol. 44, 641–644.

51. Parvez, B., and Darbar, D. (2011b). The “missing” link in atrial fibrillation heritability. J. Electrocardiol. 44.

52. Reichelt, R., Holzenburg, A., Buhle, E.L., Jarnik, M., Engel, A., and Aebi, U. (1990). Correlation between structure and mass distribution of the nuclear pore complex and of distinct pore complex components. J. Cell Biol. 110.

53. Rout, M.P., and Blobel, G. (1993). Isolation of the yeast nuclear pore complex. J. Cell Biol. 123.

54. Rout, M.P., Aitchison, J.D., Suprapto, A., Hjertaas, K., Zhao, Y., and Chait, B.T. (2000). The Yeast Nuclear Pore Complex. J. Cell Biol. 148.

55. Sachdev, R., Sieverding, C., Flötenmeyer, M., and Antonin, W. (2012). The C-terminal domain of Nup93 is essential for assembly of the structural backbone of nuclear pore complexes. Mol. Biol. Cell 23, 740–749.

56. Schwartz, M., Travesa, A., Martell, S.W., and Forbes, D.J. (2015). Analysis of the initiation of nuclear pore assembly by ectopically targeting nucleoporins to chromatin. Nucleus 6.

57. Seo, H.S., Blus, B.J., Jankovi, N.Z., and Blobel, G. (2013). Structure and nucleic acid binding activity of the nucleoporin Nup157. Proc. Natl. Acad. Sci. U. S. A. 110, 16450–16455.

58. Sjöstrand, D., Diamanti, R., Lundgren, C.A.K., Wiseman, B., and Högbom, M. (2017). A rapid expression and purification condition screening protocol for membrane protein structural biology. Protein Sci. 26, 1653–1666.

59. Sonawane, P.J., Dewangan, P., Madheshiya, P.K., Chopra, K., Kumar, M., Niranjan, S., Ansari, M.Y., Singh, J., Bawaria, S., Banerjee, M., et al. (2020). Molecular and structural analysis of central transport channel in complex with Nup93 of nuclear pore complex. Protein Sci. 29.

60. Stavru, F., Hülsmann, B.B., Spang, A., Hartmann, E., Cordes, V.C., and Görlich, D. (2006). NDC1: a crucial membrane-integral nucleoporin of metazoan nuclear pore complexes. J. Cell Biol. 173.

61. Strambio-De-Castillia, C., Niepel, M., and Rout, M.P. (2010). The nuclear pore complex: bridging nuclear transport and gene regulation. Nat. Rev. Mol. Cell Biol. 11.

62. Stuwe, T., Bley, C.J., Thierbach, K., Petrovic, S., Schilbach, S., Mayo, D.J., Perriches, T., Rundlet, E.J., Jeon, Y.E., Collins, L.N., et al. (2015). Architecture of the fungal nuclear pore inner ring complex. Science (80-.). 350.

63. Tarazón, E., Rivera, M., Roselló-Lletí, E., Molina-Navarro, M.M., Sánchez-Lázaro, I.J., España, F., Montero, J.A., Lago, F., González-Juanatey, J.R., and Portolés, M. (2012). Heart Failure Induces Significant Changes in Nuclear Pore Complex of Human Cardiomyocytes. PLoS One 7.

64. Thierbach, K., Von Appen, A., Thoms, M., Beck, M., Flemming, D., and Hurt, E. (2013). Protein interfaces of the conserved Nup84 complex from chaetomium thermophilum shown by crosslinking mass spectrometry and electron microscopy. Structure 21, 1672–1682.

65. Whittle, J.R.R., and Schwartz, T.U. (2009). Architectural Nucleoporins Nup157/170 and Nup133 Are Structurally Related and Descend from a Second Ancestral Element. J. Biol. Chem. 284.

66. Zabel, U., Doye, V., Tekotte, H., Wepf, R., Grandi, P., and Hurt, E.C. (1996). Nic96p is required for nuclear pore formation and functionally interacts with a novel nucleoporin, Nup188p. J. Cell Biol. 133.

67. Zhang, L., Tester, D.J., Lang, D., Chen, Y., Zheng, J., Gao, R., Corliss, R.F., Tang, S., Kyle, J.W., Liu, C., et al. (2016). Does Sudden Unexplained Nocturnal Death Syndrome Remain the Autopsy-Negative Disorder: A Gross, Microscopic, and Molecular Autopsy Investigation in Southern China. Mayo Clin. Proc. 91.

68. Zhang, X., Chen, S., Yoo, S., Chakrabarti, S., Zhang, T., Ke, T., Oberti, C., Yong, S.L., Fang, F., Li, L., et al. (2008). Mutation in Nuclear Pore Component NUP155 Leads to Atrial Fibrillation and Early Sudden Cardiac Death. Cell 135, 1017–1027.

69. Adams, P.D., Afonine, P. V., Bunkóczi, G., Chen, V.B., Davis, I.W., Echols, N., Headd, J.J., Hung, L.-W., Kapral, G.J., Grosse-Kunstleve, R.W., et al. (2010). PHENIX : a comprehensive Python-based system for macromolecular structure solution. Acta Crystallogr. Sect. D Biol. Crystallogr. 66.

70. Afonine, P. V., Poon, B.K., Read, R.J., Sobolev, O. V., Terwilliger, T.C., Urzhumtsev, A., and Adams, P.D. (2018). Real-space refinement in PHENIX for cryo-EM and crystallography. Acta Crystallogr. Sect. D Struct. Biol. 74.

71. Davis, I.W., Murray, L.W., Richardson, J.S., and Richardson, D.C. (2004). MOLPROBITY: structure validation and all-atom contact analysis for nucleic acids and their complexes. Nucleic Acids Res. 32.

72. Kandiah, E., Giraud, T., de Maria Antolinos, A., Dobias, F., Effantin, G., Flot, D., Hons, M., Schoehn, G., Susini, J., Svensson, O., et al. (2019). CM01: a facility for cryo-electron microscopy at the European Synchrotron. Acta Crystallogr. Sect. D Struct. Biol. 75.

73. McGuffin, L.J., Bryson, K., and Jones, D.T. (2000). The PSIPRED protein structure prediction server. Bioinformatics 16.

74. Pettersen, E.F., Goddard, T.D., Huang, C.C., Couch, G.S., Greenblatt, D.M., Meng, E.C., and Ferrin, T.E. (2004). UCSF Chimera?A visualization system for exploratory research and analysis. J. Comput. Chem. 25.

75. Punjani, A., Rubinstein, J.L., Fleet, D.J., and Brubaker, M.A. (2017). cryoSPARC: algorithms for rapid unsupervised cryo-EM structure determination. Nat. Methods 14.

76. Scheres, S.H.W. (2012). RELION: Implementation of a Bayesian approach to cryo-EM structure determination. J. Struct. Biol. 180.

77. Schwede, T. (2003). SWISS-MODEL: an automated protein homology-modeling server. Nucleic Acids Res. 31.

78. Zhang, K. (2016). Gctf: Real-time CTF determination and correction. J. Struct. Biol. 193.

79. Zheng, S.Q., Palovcak, E., Armache, J.-P., Verba, K.A., Cheng, Y., and Agard, D.A. (2017). MotionCor2: anisotropic correction of beam-induced motion for improved cryo-electron microscopy. Nat. Methods 14.

